# Distinct roles of BRCA2 in replication fork protection in response to hydroxyurea and DNA interstrand crosslinks

**DOI:** 10.1101/811968

**Authors:** Kimberly A. Rickman, Ray Noonan, Francis P. Lach, Sunandini Sridhar, Anderson T. Wang, Avinash Abhyankar, Michael Kelly, Arleen D. Auerbach, Agata Smogorzewska

## Abstract

DNA interstrand crosslinks (ICLs) are a form of DNA damage that requires the interplay of a number of repair proteins including those of the Fanconi anemia (FA) and the homologous recombination (HR) pathways. Pathogenic variants in the essential gene *BRCA2/FANCD1,* when monoallelic, predispose to breast and ovarian cancer, and when biallelic, results in a severe subtype of Fanconi anemia. BRCA2 function in the FA pathway is attributed to its role as a mediator of the RAD51 recombinase in HR repair of the programmed DNA double strand breaks (DSB). BRCA2 and RAD51 functions are also required to protect stalled replication forks from nucleolytic degradation during response to hydroxyurea (HU). While RAD51 has been shown to be necessary in the early steps of ICL repair to prevent aberrant nuclease resection, the role of BRCA2 in this process has not been described. Here, based on the analysis of BRCA2 DNA binding domain (DBD) mutants discovered in FA patients presenting with atypical FA-like phenotypes, we establish that BRCA2 is necessary for protection of DNA at an ICL. Cells carrying DBD BRCA2 mutations are sensitive to ICL inducing agents but resistant to HU treatment consistent with relatively high HR repair in these cells. BRCA2 function at an ICL protects against DNA2-WRN nuclease-helicase complex and not the MRE11 nuclease implicated in the resection of HU-stalled replication forks. Our results also indicate that unlike the processing at HU-stalled forks, function of the SNF2 translocases (SMARCAL1, ZRANB3, or HLTF), implicated in fork reversal, are not an integral component of the ICL repair, pointing to a different mechanism of fork protection at different DNA lesions.

## Introduction

DNA interstrand crosslinks (ICLs) are a deleterious form of DNA damage that covalently link the Watson and Crick strands of DNA. ICLs can be produced by exogenous compounds such as mitomycin C (MMC), diepoxybutane (DEB), cisplatin, psoralen, and nitrogen mustards or by naturally occurring biological metabolites such as aldehydes [1–3].

The importance of the proper repair of ICLs is emphasized by the rare genetic disorder, Fanconi anemia (FA). FA is characterized by developmental abnormalities, bone marrow failure (BMF), predisposition to solid tumors and leukemia, and cellular hypersensitivity to crosslinking agents [4]. FA results from pathogenic variants in one of the 22 *FANC* genes (FANCA-W) whose protein products are required for proper ICL repair [2, 5–7].

When an ICL is encountered during DNA replication it causes fork stalling and FA pathway activation [8, 9]. The removal of an ICL is a multistep process requiring activation of the FA core complex and monoubiqutination of FANCD2 and FANCI [8, 10, 11]. Monoubiquitinated FANCD2 and FANCI form a heterodimer that is recruited to chromatin and is required for ICL processing, which entails nucleolytic unhooking of the crosslinked DNA [12–15]. Unhooking of the ICL enables translesion bypass on one strand and double strand break (DSB) repair by homologous recombination (HR) on the second strand [16–19].

A number of FA proteins, BRCA2/FANCD1, PALB2/FANCN, FANCJ/BRIP1, RAD51C/FANCO, RAD51/FANCR, and BRCA1/FANCS, are known for facilitating HR [16-18, 20-22]. *BRCA2/FANCD1* is an essential gene and single allele pathogenic variants predispose to breast and ovarian cancer and biallelic pathogenic variants result in a subtype of Fanconi anemia, FA-D1 [16]. FA is a heterogeneous disease, but even within the disease spectrum, patients with biallelic pathogenic variants in *BRCA2/FANCD1* are phenotypically distinct from the most common complementation groups, FA-A, FA-C, and FA-G. A higher proportion of FA-D1 patients have developmental abnormalities and nearly one hundred percent have a malignancy by five years of age [23], which is most likely due to HR deficiency.

Functional analysis of BRCA2 has largely focused on canonical HR and the role of BRCA2 in ICL repair has been associated with the repair of DSBs generated by programmed incisions at the ICL. Outside of their role in HR and ICL repair, BRCA2 and RAD51 along with a number of other recently described proteins, function in replication fork protection [24]. In the absence of replication fork protection, newly synthesized DNA is degraded at hydroxyurea (HU) stalled replication forks and a number of nucleases including MRE11, CTIP, and EXO1 have been implicated in the process [24–26].

Another nuclease, DNA2, has also been shown to resect DNA at ICLs in cells expressing the RAD51/FANCR separation of function mutant, p.T131P, identified in an individual with FA-like syndrome. The mutant RAD51 p.T131P has a dominant negative effect on RAD51 function that does not seem to affect HR at cellular levels but disrupts the function of RAD51 at ICLs, suggesting a fork protection role in ICL repair. The requirement for BRCA2 in the early steps of ICL repair to prevent aberrant resection has not previously been determined. Here we investigated the requirements of BRCA2 with RAD51 in fork protection of ICLs and demonstrate the two proteins are both required to prevent hyper-resection by the DNA2-WRN nuclease-helicase complex, but not MRE11. These studies were performed using BRCA2 DNA binding domain (DBD) mutants discovered in FA patients and these variants were determined to confer loss of replication fork protection but only moderate HR deficiency. Our results indicate that the BRCA2 DBD is required for replication fork protection and that BRCA2 fork protection at HU-stalled forks and ICLs are distinct processes.

## Results

### Atypical presentation of Fanconi anemia in individuals with biallelic *BRC*A2/*FANCD1* DNA binding domain variants

Two female siblings, enrolled in the International Fanconi Anemia Registry (IFAR), with unknown causative gene mutations, were born with a multitude of congenital abnormalities and had mildly elevated levels of chromosomal breakage at birth (see **Table S1** for clinical presentation). Biallelic *BRCA2*/*FANCD1* variants (c.2330dupA and c.8524C>T) were identified by Whole Exome Sequencing (WES) and no other likely-pathogenic FA gene variants were observed. These results were surprising since neither sibling displayed the typical clinical findings of the FA-D1 complementation group, with no history of malignancy or bone marrow failure at the ages of 20 and 23. There is no reported family history of FA, but there are cases of breast cancer that were diagnosed later in life (above 60 years of age), individuals with skin cancer in the family, and early onset colorectal cancer in the father (40 years-old) (**Figure 1A**).

**Figure 1:**
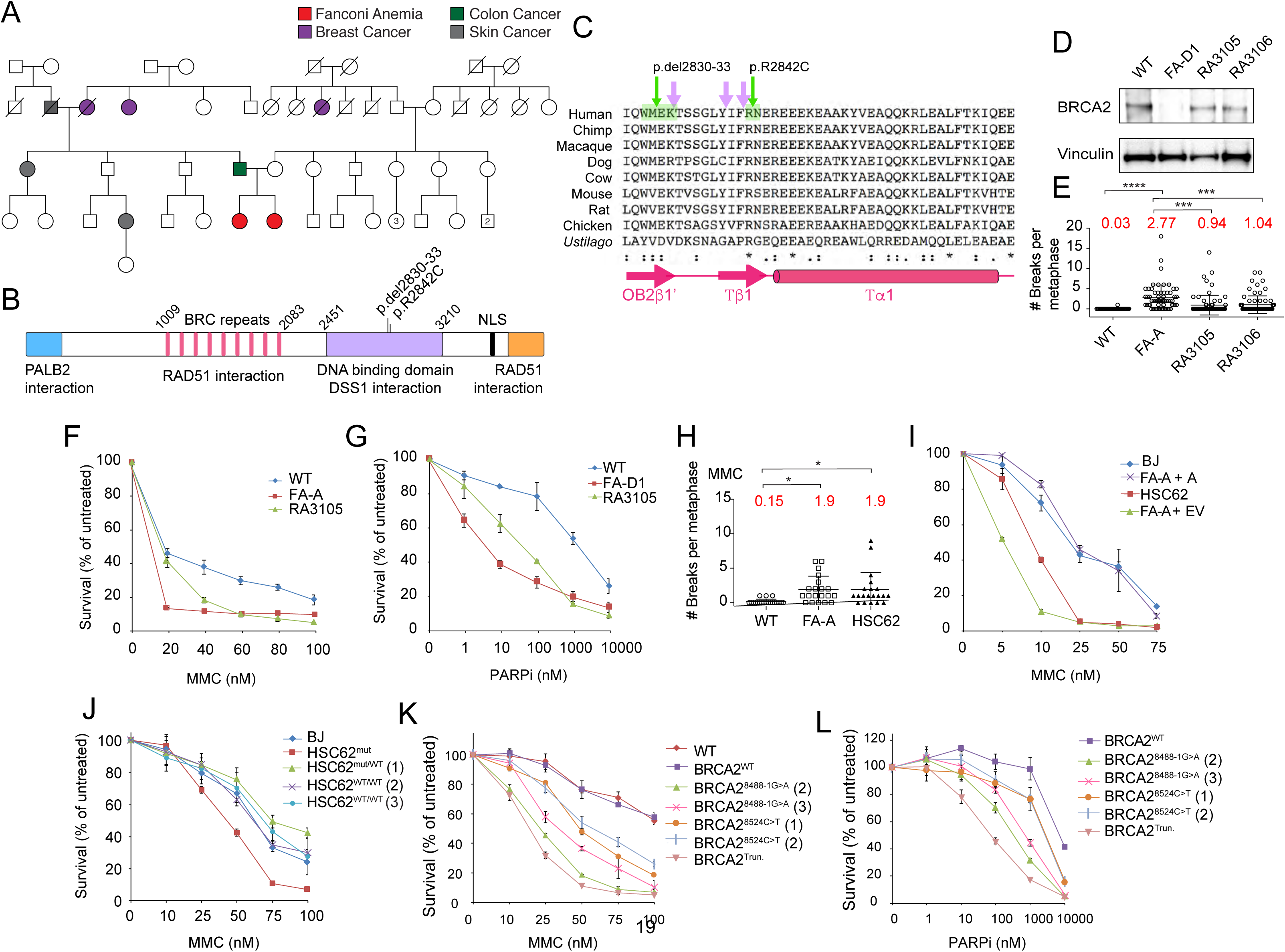
BRCA2 variants identified in individuals with atypical Fanconi anemia. (A) Family pedigree showing a sibling pair with Fanconi anemia (red circles) who are compound heterozygous for *BRCA2* variants c.2330dupA (maternal inheritance) and c.8524C>T (paternal inheritance). Family history of breast cancer (purple, all diagnosed in 60s and 70s), skin cancer (grey), and colon cancer (green, diagnosed at 40 years old). (B) Schematic of BRCA2 domain structure and key interacting proteins (C) Alignment of exon 20 BRCA2 DBD peptide sequence demonstrating it is evolutionary conserved across many species. In green are the aa residues modified by the patient variants, p.W2830_K2833del (c.8488-1G>A) and p.R2842C (c.8524C>T). Purple arrows indicate AA residues that contact DNA [63]. (D) Immunoblot showing BRCA2 levels in WT (RA2985) control, FA-D1 (RA2525), and patient RA3105 and RA3106 LCLs. (E) Quantification of chromosome breaks following DEB treatment of WT (RA2985), FA-A (RA2939), and patient RA3105 and RA3106 LCLs. (F-G) Cell survival assays of patient derived lymphoblast cell lines (LCLs) RA3105, FA-A (RA2939), WT (RA2985), and FA-D1 (RA2525) after MMC and PARP inhibitor olaparib (PARPi) treatment. Relative cell survival was normalized to untreated controls to give percent survival. Error bars indicate s.d. (H) Quantification of chromosome breaks following MMC treatment of BJ wild type fibroblasts, FA-A patient fibroblasts, and HSC62 fibroblasts. (I) Cell survival of HSC62 (c.8488-1G>A) fibroblasts compared to BJ WT fibroblast, FA-A patient fibroblast, complemented FA-A patient cells (RA3087) expressing wild type *FANCA* (FA-A+A) or empty vector (FA-A+EV). Cells were treated with increasing concentrations of MMC. Relative cell survival was normalized to untreated controls to give the percent survival. Error bars indicate s.d. (J) Cell survival of MMC treated HSC62 uncorrected patient cell line (HSC62^mut^) compared to BJ WT fibroblast and CRISPR/Cas9 corrected wild type HSC62 (HSC62^WT^) clones 1-3. (K-L) Cell survival of BJ WT fibroblasts, and CRISPR/Cas9 targeted BJ fibroblasts: BJ WT fibroblast clone (BRCA2^WT^), c.8488-1G>A BJ clones (BRCA2^8488-1G>A^), c.8524C>T BJ clones (BRCA2^8524C>T)^, and exon 20 *BRCA2* frameshift mutant (BRCA2^Trun.^). Cells were treated with increasing concentrations of MMC or PARPi. Error bars indicate s.d.

The frameshift c.2330dupA variant of exon 11 (maternal origin) results in premature truncation of BRCA2 (p.Asp777Glufs*11) and has previously been described in hereditary breast and ovarian cancer (HBOC) (Figure S1A). The c.8524C>T missense variant of exon 20 (paternal origin) results in an p.Arg2842Cys residue change in the highly conserved DNA binding domain (DBD) of BRCA2 and has previously been identified as a variant of unknown significance (VUS) in HBOC (**Figure 1B-C,** Figure S1B-C). At the protein level, the missense variant results in the p.Arg2842Cys change at a highly conserved residue at the base of the BRCA2 Tower domain of the DBD (Figure S1C). Sequencing of peripheral blood and lymphocytes demonstrated the presence of both variants and no evidence of somatic mosaicism.

A third individual with FA, biallelic *BRCA2* variants, and an atypical presentation, was identified in the literature [16]. This individual was homozygous for the c.8488-1G>A variant (alias “IVS19-1G>A”) that alters the splice acceptor site of exon 20. cDNA analysis demonstrated the use of an alternate splice acceptor that results in the loss of 12 base pairs (bp) of exon 20 and translates into p.Trp2830_Lys2833del [16] (**Figure 1B-C**). Amino acid residues 2830-2833 are located within the DBD at the transition of the OB2 fold and the base of the Tower domain (**Figure 1B-C, Figure S1C**). This individual was 30 years of age at last follow up, was born with a thumb malformation, but had no history of bone marrow failure or malignancy. Similar to the sibling pair, chromosomal breakage was modest [16].

### *BRCA*2 DNA binding domain variants identified in FA patients confer defects in the response to replication stress

Lymphoblastoid cell lines (LCL) (RA3105 and RA3106) were derived from the sibling pair with compound heterozygous *BRCA2* variants, c.2330dupA and c.8524C>T. FA pathway activation, monitored by FANCI ubiquitination, was normal in patient-derived LCLs (Figure S1D). Analysis of BRCA2 expression by western blot demonstrated a full length (∼390 kDa) band, the presumed product of the c.8524C>T allele, for both patient cell lines (**Figure 1D**). DEB-induced breakage analysis confirmed previous clinical data that breakage was elevated, but not to levels of the typical FANCA deficient (FA-A) LCLs (RA2939) (**Figure 1E**). RA3105 LCL displayed hypersensitivity to the crosslinking agents MMC and DEB, but to a lesser degree than RA2939 (**Figure 1F,** Figure S1F). RA3105 was also hypersensitive to replication stress inducing agents including olaparib, a PARP inhibitor (PARPi), and camptothecin (CPT), a topoisomerase I inhibitor (**Figure 1G,** Figure S1G).

Similarly, analysis of patient-derived fibroblasts, HSC62, [16] from the individual with homozygous c.8488-1G>A variant also revealed more moderate chromosomal breakage to DEB and MMC and cellular hypersensitivity to crosslinking agents (**Figure 1H**, I, Figure S1O). Interestingly, the cells were not hypersensitive to ionizing radiation (IR), but were sensitive to replication stress induced by CPT and PARPi (Figure S1J-L). In contrast, the cells were not sensitive to replication stress produced by the agents aphidicolin and HU (Figure S1M-N).

We complemented the HSC62 patient fibroblast cell line to demonstrate that the c.8488-1G>A variant caused the observed defects. The homozygous c.8488-1G>A variant was corrected to wild type at the endogenous locus using CRISPR/Cas9 gene targeting. Both heterozygous and homozygous clones were recovered (HSC62^WT/MUT^ or HSC62^WT/WT^) (Figure S1 P). cDNA analysis demonstrated that restoration of the splice acceptor base (A>G) in HSC62^WT/MUT^ or HSC62^WT/WT^ clones restored the cDNA exon 19-20 junction (**Figure S**1 Q). Both HSC62^WT/MUT^ and HSC62^WT/WT^ clones rescued hypersensitivity to replication stress inducing agents MMC, CPT, and PARPi **(Figure 1J and** S1S-T).

For a direct comparison of the *BRCA2* DNA binding domain variants, we generated isogenic cell lines by introducing the variants, c.8524C>T (p.R2842C) and c.8488-1G>A (p.Trp2830_Lys2833del), into wild type BJ fibroblasts with CRISPR/Cas9 gene editing (Figure S2A-B). Knock-in of the *BRCA2* c.8488-1G>A variant in BJ fibroblasts conferred the same splicing defect observed in HSC62 cells (Figure S2A). Western blot analysis of BRCA2 demonstrated a ∼390 kDa band for all mutants except for *BRCA2* clones containing exon 20 frameshift variants obtained in parallel using CRISPR/Cas9 gene targeting (Figure S2C). The *BRCA2* frameshift mutant is homozygous c.8531dupA with a predicted p.R2845Kfs*22 truncation (BRCA2^Trun.^). Analysis of cellular sensitivity of the BRCA2 DBD mutants revealed that presence of both DBD variants sensitize cells to MMC, PARPi, CPT but not aphidicolin, recapitulating phenotypes of patient HSC62 fibroblasts (**Figure 1K-L,** S2 D-E).

**Figure 2:**
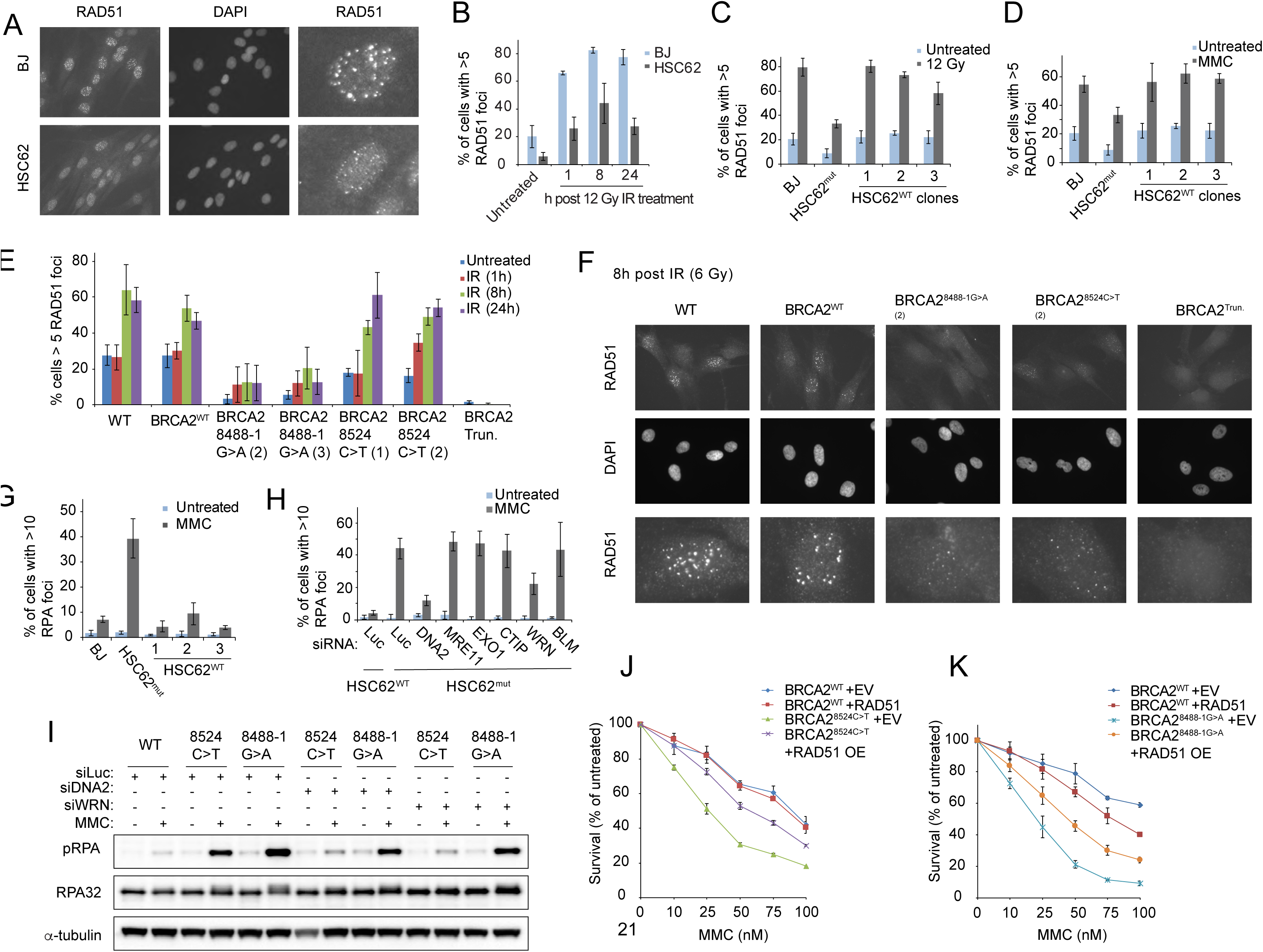
Defective ICL repair in BRCA2 DBD mutants results in hyperactivation of RPA that is WNR and DNA2 dependent. (A) Immunofluorescence images of RAD51 foci, 8h following 12 Gy ionizing radiation (IR) of BJ WT fibroblast and patient derived HSC62 fibroblast, detected with anti-RAD51 antibody. Third row images are individual cells enlarged to better demonstrate differences in RAD51 foci size. (B) Quantification of RAD51 foci 1h, 8h, and 24h following 12 Gy IR of BJ WT fibroblast and HSC62 fibroblast. Error bars indicate s.d. of two independent experiments (≥200 cells per experiment). (C) Quantification of RAD51 foci 8h after 12 Gy IR of BJ WT fibroblast, wild type HSC62 - (HSC62^WT^) clones 1-3, and HSC62 uncorrected patient cell line (HSC62^mut^). (D) Quantification of RAD51 foci 24h following 1h treatment with 3 µM MMC. Error bars indicate s.d. of three independent experiments (≥200 cells per experiment). (E) Quantification of RAD51 foci in isogenic BJ fibroblasts clones at 1h, 8h and 24h following 6 Gy IR of BJ WT fibroblasts, BJ WT fibroblast clone (BRCA2^WT^), BRCA2^8488-1G>A^ BJ clones 2-3, BRCA2^8524C>T^ BJ clones 1-2, and a *BRCA2* homozygous truncation mutant, c.8531dupA (BRCA2^Trun^). Error bars indicate s.d. of three independent experiments (≥200 cells per experiment) (F) Representative images of RAD51 foci in isogenic BJ fibroblasts clones, 8h post 6 Gy IR, detected by immunofluorescence with anti-RAD51 antibody. Third row images are individual cells enlarged to better demonstrate differences in RAD51 foci size. (G) Quantification of RPA foci 24h following 1h treatment with 3 μM MMC of BJ WT fibroblast, CRISPR/Cas9 corrected wild type HSC62 clones (HSC62^WT^), and HSC62 uncorrected patient cell line (HSC62^mut^). (H) Quantification of RPA foci 24h following 1h treatment with 3 μM MMC in HSC62^mut^ cells depleted of DNA2, MRE11, EXO1, CTIP, WRN, or BLM by siRNA compared to luciferase control (Luc). Error bars indicate s.d. of four independent experiments. (I) Immunoblot analysis of RPA phosphorylation in isogenic BJ fibroblasts clones 24h post 1h treatment with 3 μM MMC. BRCA2^WT^, BRCA2^8524C>T^, and BRCA2^8488-1G>A^ BJ fibroblast cells were transfected with siRNA control luciferase (Luc) or siRNAs targeting DNA2 or WRN. (J-K) MMC cell survival of BJ BRCA2^WT^, BRCA2^8488-1G>A^, and BRCA2^8524C>T^ fibroblasts overexpressing (OE) WT RAD51 or empty vector (EV) control. Relative cell survival was normalized to untreated controls to give percent survival. Error bars indicate s.d.

### *BRCA*2 DNA binding domain variants confer defects in RAD51 recruitment to ssDNA after IR and MMC

To determine the impact of DBD variants on the ability of BRCA2 to load RAD51 onto ssDNA following DNA damage, we analyzed RAD51 foci formation after IR and MMC. Levels of RAD51 foci and foci size were reduced after IR and MMC treatment in HSC62 cells, which was rescued by complementation by CRISPR/Cas9 gene editing **(Figure 2A-D,** Figures S2F-G). Analysis of isogenic BJ cell lines with DBD mutations also demonstrated defects in RAD51 foci formation following IR and MMC **(Figure 2E-F,** Figure S2H-I). The c.8488-1G>A variant had a stronger impact on RAD51 foci formation, resulting in fewer cells with RAD51 foci and reduced foci size. The c.8524C>T mutant did not show a significant reduction in the number of cells with RAD51 foci; however, the foci were smaller in size **(Figure 2F,** Figures S2I). By comparison the BRCA2^Trun.^ mutant had complete loss of observable RAD51 foci. These data indicate that the BRCA2 DBD mutants are hypomorphic in their mediator function.

### Increased RPA activation in BRCA2 DBD variants is dependent on DNA2 and WRN

The previously described RAD51/FANCR p.T131P patient-derived cell line that is proficient for HR but defective in ICL repair displays hyperactivation of RPA upon MMC treatment [27]. Given that the interaction of BRCA2 and RAD51 is required for their canonical function in HR and their non-canonical function in replication fork protection at HU-stalled forks, we investigated whether BRCA2 also functions in preventing increased ssDNA generation at ICLs [27–30]. We observed an increase in RPA foci formation and RPA phosphorylation in HSC62^MUT^ cells compared to wild type fibroblasts upon MMC treatment (**Figure 2G,** Figure S3A). Similar to RAD51/FANCR p.T131P expressing patient cells, the increased RPA foci formation in HSC62 cells was also dependent on DNA2 and WRN activity, but not MRE11, EXO1, CTIP, or BLM (**Figure 2H,** Figure S3B-C). Increased RPA foci following MMC was also observed for c.8524C>T and c.8488-1G>A mutants, with a greater increase of RPA foci and activation in the c.8488-1G>A mutants **(Figure 2I,** Figure S3D-H). These results suggest that BRCA2 is functioning with RAD51 to protect against aberrant processing by DNA2 and WRN at ICLs, but not against the other effectors of DSB end resection such as MRE11, EXO1, or CTIP. Overexpression of RAD51 in the *BRCA2* c.8524C>T and c.8488-1G>A mutants partially rescued cellular sensitivity to MMC and RPA foci formation after MMC (**Figure 2J-K,** Figure S3I-J). This data supports that RAD51 and BRCA2 function interdependently in their roles at ICLs.

To determine if blocking ICL unhooking or nuclease mediated fork collapse [13, 14, 31-33] would rescue RPA foci formation in *BRCA2* DBD mutant cells following MMC, we depleted SLX4 and MUS81. Depletion of either SLX4 or MUS81 did not rescue RPA hyperactivation in the *BRCA2^8524C>T^ and BRCA2^8488-1G>A^* cells (Figure S3K-M). SLX4 depletion further increased RPA activation and foci formation, indicating further defects in ICL repair in its absence, which may be the result of loss of the function of the associated nucleases.

### ICLs are a substrate of nucleolytic processing in the absence of a functioning FA pathway

Having demonstrated that BRCA2 and RAD51 share a role in protecting ICLs from over-resection by DNA2 and WRN, we investigated whether other FA proteins are also required for protection against DNA hyper-resection at ICLs. Analysis of a panel of FA patient-derived cells with mutations in FANCA, FANCL, FANCD2, FANCI, FANCJ, and SLX4/FANCP demonstrated increased RPA foci formation following MMC treatment for all complementation groups **(Figure 3A)**. To determine if the genetic requirement for RPA suppression was the same as in BRCA2 and RAD51 mutant cells, DNA2 and WRN were depleted in a complemented pair of FANCA patient-derived cells **(Figure 3B-C,** Figure S3N-O). Interestingly, the dependence on DNA2 was the same, but the helicase dependency was different, as WRN did not rescue RPA levels but BLM depletion did **(Figure 3C,** Figure S3P-Q). These data demonstrate a dependence on the FA core complex and pathway associated proteins to prevent resection of ICLs by DNA2 and BLM. They also suggest that different nuclease-helicase pairs engage when ICL repair is halted at different stages of the process.

**Figure 3:**
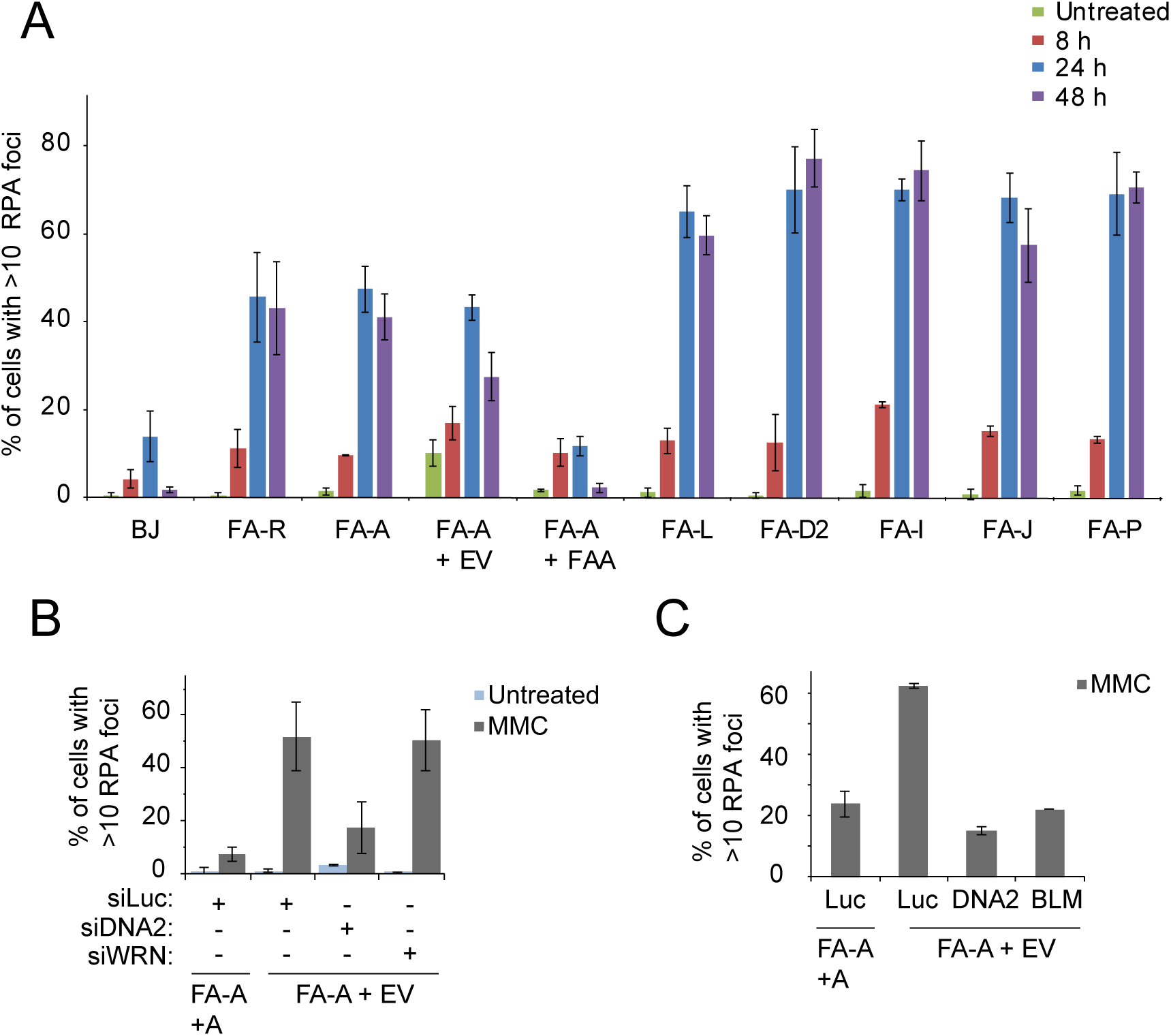
Proper ICL repair is required to prevent aberrant nuclease processing. (A) Quantification of RPA foci 8h, 24h and 48h following 1h treatment with 3 μM MMC of FA patient derived fibroblasts compared to BJ wild type fibroblasts. Patient cells lines from FA complementation group FA-R (RAD51/FANCR) (FA-R^mut^), FA-A (FANCA) (FA-A^mut^), FA-L (FANCL) (FA-L^mut^), FA-D2 (FANCD2) (FA-D2^mut^), FA-I (FANCI) (FA-I^mut^), FA-J (FANCJ) (FA-J^mut^), and FA-P (SLX4/FANCP) (FA-P^mut^). FA-A patient complemented cell lines were generated by transducing WT *FANCA* cDNA or EV. Error bars indicate s.d. of two independent experiments. (B) FA-A patient cells expressing WT *FANCA* (FA-A+A) or empty vector (FA-A+EV) were transfected with siRNA control luciferase (Luc) or siRNAs targeting DNA2 and WRN. Quantification of RPA foci 24h following 1h treatment with 3 μM MMC. Error bars indicate s.d. of two independent experiments. (C) FA-A+EV were transfected with siRNA Luc or siRNAs targeting DNA2 and BLM. Quantification of RPA foci 24h following 1h treatment with 3 μM MMC. Error bars indicate s.d. of two independent experiments.

## Determination of homologous recombination efficiency in DNA binding domain mutants

To determine the HR proficiency of *BRCA2^8488-1G>A^*and *BRCA2^8524C>T^* cells, we utilized a HDR assay that targets DSBs at the LMNA locus [34, 35]. The assay was performed in HEK293T cells after CRISPR-Cas9 gene editing to have either BRCA2 DBD variants or the exon 27 p.S3291A variant, previously reported to have an effect on replication fork protection but not on HR **(Figure S4A-B)** [28, 36]. Compared to wild type cells, HR in all BRCA2 clones, including the S3291A mutant, was moderately decreased **(Figure 4A).** Cells with DBD *BRCA2^8488-1G>A^*and *BRCA2^8524C>T^* variants showed similar decreases in HR levels to approximately half that of wild type cells but retained significantly more HR activity than cells depleted of *RAD51* and *BRCA2* or *BRCA2^Trun^*cells. Two of the c.8488-1G>A mutants appeared to express lower BRCA2 levels (Figure S4B), suggesting these may be hemizygous clones, but this did not further impair HR levels when compared to similar HR levels in a third homozygous clone.

**Figure 4:**
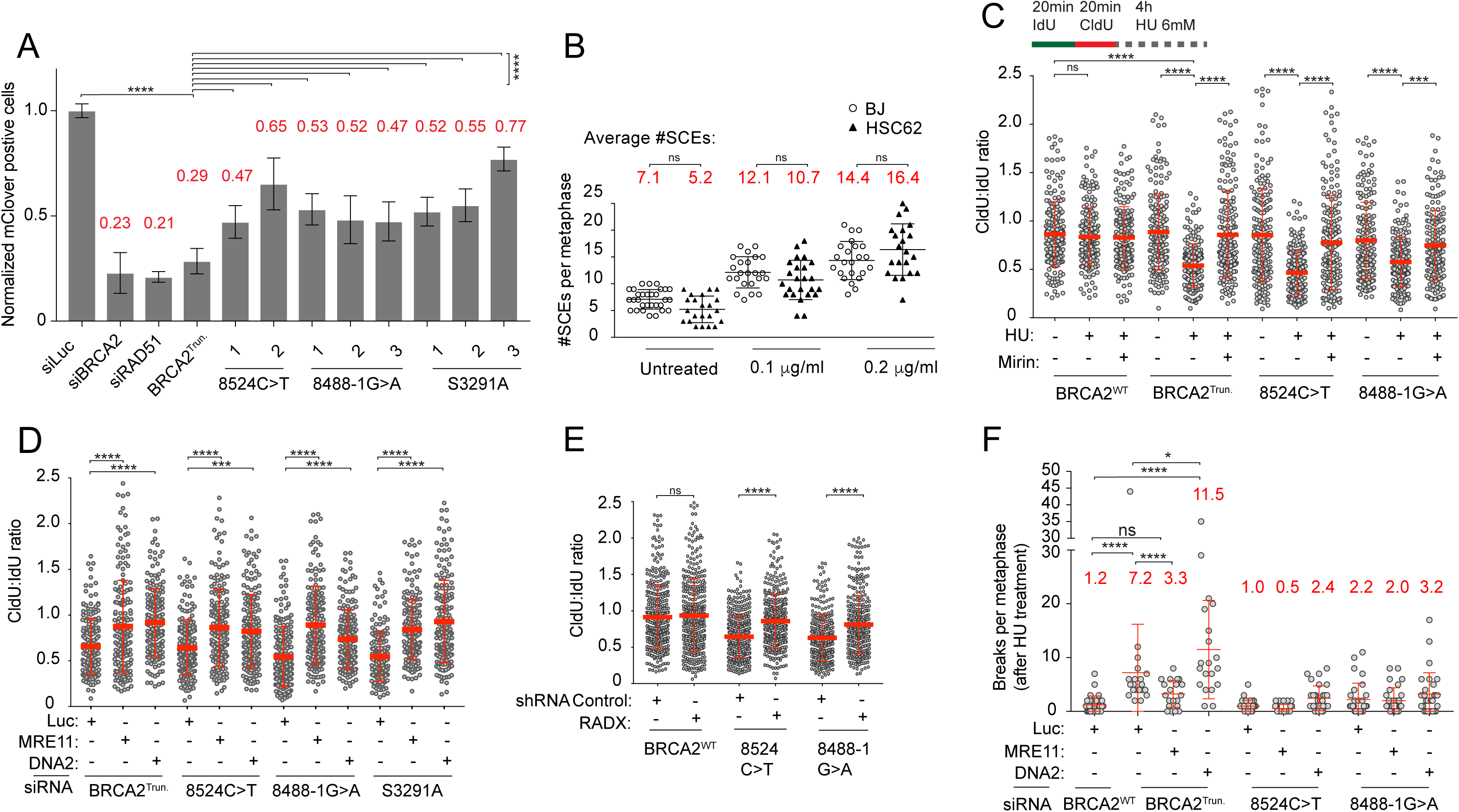
*BRCA2* DBD and C-terminal domain variants confer a moderate defect in HR and disrupt replication fork protection function. (A) Levels of mClover positive cells were normalized to WT HEK293T (siLuc). Error bars indicate s.d. of three independent experiments performed in triplicate. (B) Sister chromatid exchange (SCE) assay in BJ WT fibroblast and HSC62 patient derived fibroblast following treatment with MMC (0.1 μg/ml or 0.2 μg/ml). (C) Isogenic BJ fibroblast BRCA2 mutants, BRCA2^Trun.^, BRCA2^8524C>T^ and BRCA2^8488-1G>A^, were analyzed for replication fork resection. Cells were labeled with DNA analogs, IdU for 20 minutes and then CldU for 20 minutes. Cells were then incubated in 6 mM HU with and without MRE11 inhibitor mirin (50 uM) for 4h before being harvested. DNA fibers were prepared and visualized by immunofluorescence detection of IdU and CldU and measured. Error bars indicate s.d. (D) Isogenic BJ fibroblast BRCA2 mutants, BRCA2^Trun.^, BRCA2^8524C>T^, BRCA2^8488-1G>A^, and BRCA2^S3291A^, were transfected with siRNA control luciferase (Luc) or siRNAs targeting DNA2 or MRE11. Cells were treated and labeled with DNA analogs as above. Error bars indicate s.d. (E) BJ fibroblast with BRCA2 variants, BRCA2^8524C>T^ and BRCA2^8488-1G>A^, were analyzed for replication fork resection when depleted of RADX by shRNA or transduced with shRNA control (shCONT.). Cells were treated and labeled with DNA analogs as above. Data of two replicated plotted. Error bars indicate s.d. (F) Quantification of chromosome breaks in isogenic BJ fibroblast *BRCA2* mutants following 5h of 6 mM HU and released into colcemid. Breakage was not significantly increased in BRCA2^8524C>T^ and BRCA2^8488-1G>A^ compared to BRCA2^WT^.

Given the normal resistance to IR in HSC62 fibroblasts, we assessed sister chromatid exchange (SCEs) levels as a readout of HR [37]. SCEs were induced by increasing concentrations of MMC or depletion of BLM. There was no significant difference in SCE levels observed in wild type BJ fibroblasts and HSC62 cells **(Figure 4B,** Figure S4E-G); however, SCE levels were suppressed in *BRCA2^Trun^*fibroblasts (Figure S4H). These observations suggest that the DNA binding domain defect in HSC62 cells, while decreasing RAD51 foci formation, does not significantly reduce HR as observed by normal resistance to IR and SCE levels in these cells. Taken together, the variants moderately reduce HR at Cas-9 targeted DSBs but do not impact cellular HR readouts, which is similar to the behavior of cells carrying the RAD51 p.T131P mutation [27].

## The BRCA2 DNA binding domain is required for replication fork protection at HU-stalled forks

To determine the requirement for the BRCA2 DBD in replication fork protection after HU treatment, *BRCA2^8524C>T^ and BRCA2^8488-1G>A^* cells were examined by DNA fiber analysis. Replication fork protection by BRCA2 has largely been attributed to the C-terminal RAD51 interacting domain by analysis of the BRCA2 p.S3291A variant [28]. Analysis of BRCA2^Trun.^, *BRCA2^8524C>T^,* and *BRCA2^8488-1G>A^*cells demonstrated defects in replication fork protection of HU-stalled forks as measured by the degradation of nascent DNA tracks labeled with nucleotide analogs, IdU and CldU. As previously reported, nascent strand degradation in the absence of BRCA2 was rescued by the MRE11 inhibitor mirin and MRE11 depletion **(Figure 4C-D)**. These data demonstrate that the *BRCA2^Trun.^, BRCA2^8524C>T^,* and *BRCA2^8488-1G>A^* cells are all similarly defective for replication fork protection and that the DBD is required for protection of replication forks from MRE11 processing. Depletion of DNA2 also rescues resection after HU in all of the BRCA2 mutants including *BRCA2^Trun.^*, *BRCA2^8524C>T^, BRCA2^8488-1G>A^*, and *BRCA2^S3291A^* (**Figure 4D)**. RADX depletion has been shown to rescue nascent strand degradation at HU-stalled replication forks in BRCA2 deficient cells without restoring HR function [38]. Consistent with these studies, depletion of RADX in the BRCA2 DBD mutant cells did not rescue HR defects (Supplemental Figure SF5A-D) but did rescue nascent strand degradation **(Figure 4E)**. Taken together, these data demonstrate that both the DBD and C-terminal domain of BRCA2 are required for proper replication fork protection at HU-stalled forks, and that both domains are required to protect against degradation by the nucleases MRE11 and DNA2.

Although all of the BRCA2 mutants showed similar levels of nascent strand resection as measured by DNA fibers, the levels of chromosomal breakage differed **(Figure 4F,** Figures S5 E-F). Metaphases were analyzed after 5 hours of 6 mM HU and release into colcemid. *BRCA2^Trun.^*cells showed a large increase in genomic instability upon stalling with HU in comparison to WT and the other *BRCA2* mutants. Cells with *BRCA2^8524C>T^*and *BRCA2^S3291A^* variants did not show an elevation in breakage and *BRCA2^8488-1G>A^*cells had a mild increase. The elevated chromosomal breakage in *BRCA2^Trun.^* cells were reduced by MRE11 depletion, but exacerbated by DNA2 depletion **(Figure 4F,** Figure S5E). DNA2 depletion resulted in a mild increase in breakage for all mutants but resulted in a synergistic increase in *BRCA2^Trun^.* Previous studies have reported elevated breakage resulting from replication fork degradation in p.S3291A expressing cells and BRCA2 deficient cells [28, 29]. In contrast, the newly characterized *BRCA2^8524C>T^* or *BRCA2^S3291A^*cells in this study do not have a significant increase in breakage after HU, despite having levels of fork degradation similar to *BRCA2^Trun^* **(Figure 4F,** Figure S5F). Our data demonstrate that different levels of BRCA2 function impairment have different consequences on HU-stalled forks and that replication fork resection at HU-stalled forks does not always manifest in chromosomal breakage. How this breakage occurs in *BRCA2* depleted or LOF cells needs to be investigated further, but like nascent DNA degradation, it is partially dependent on MRE11.

**Figure 5:**
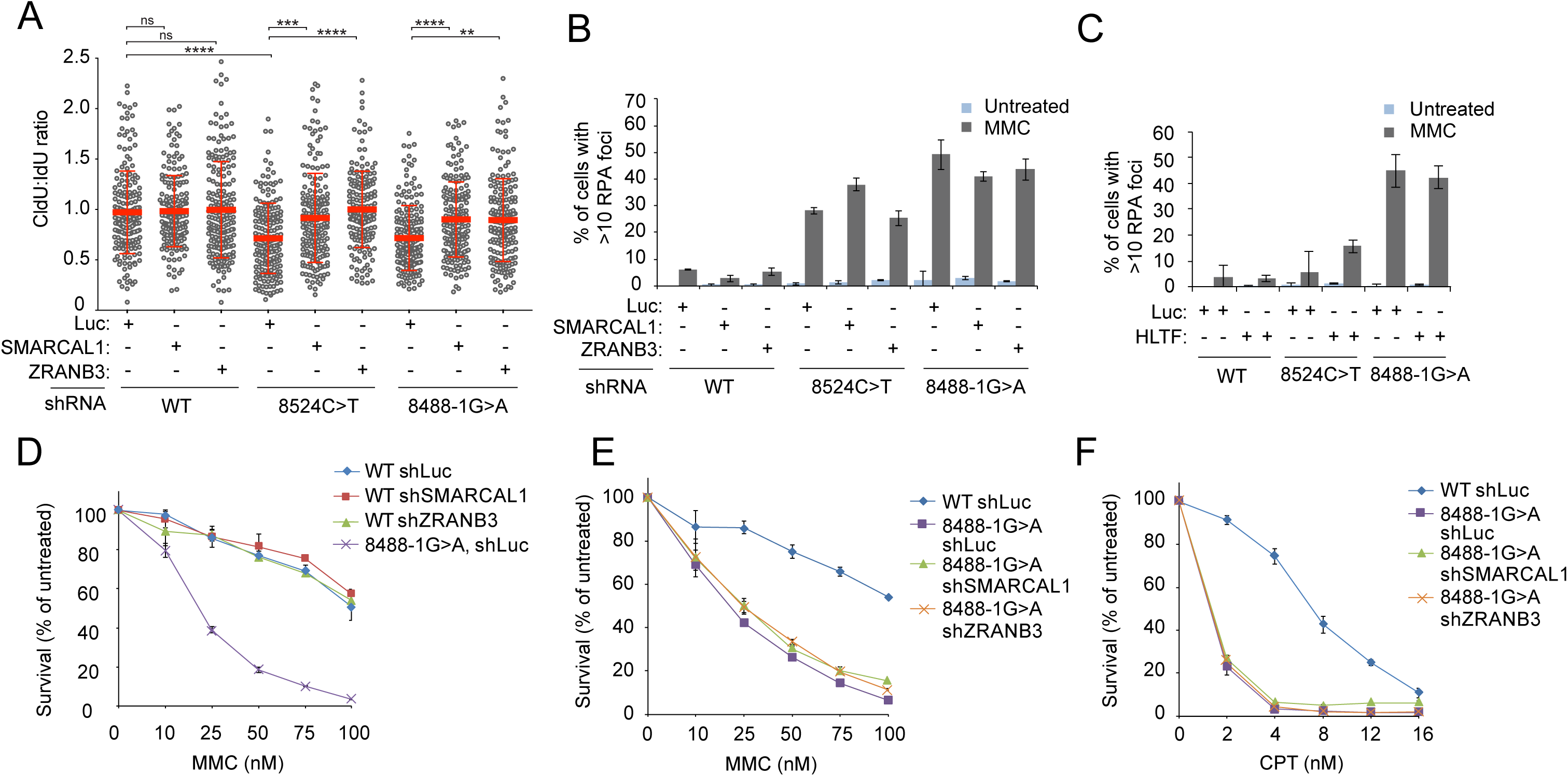
SNF2 translocases are not required for ICL repair. (A) BJ fibroblast mutants BRCA2^8524C>T^ and BRCA2^8488-1G>A^, were analyzed for replication fork resection when depleted of either SMARCAL1 or ZRANB3 by shRNA or transduced with control shRNA (shLuc). Cells were labeled with DNA analogs, IdU for 20 minutes and then CldU for 20 minutes. Cells were then incubated in 6 mM HU for 4h before being harvested. DNA fibers were prepared and visualized by immunofluorescence detection of IdU and CldU and measured. Error bars indicate s.d. (B) Quantification of RPA foci in isogenic BJ fibroblasts clones 24h following 1h treatment with 3 µM MMC in cells depleted of SMARCAL1 or ZRANB3. Error bars indicate s.d of two independent experiments. (C) Quantification of RPA foci in BJ fibroblasts clones 24h following 1h treatment with 3 uM MMC in cells depleted of HLTF. Error bars indicate s.d of two independent experiments. (D) MMC cell survival of isogenic BJ BRCA2^WT^ fibroblasts depleted of SMARCAL1 or ZRANB3 by shRNA or transduced with shRNA luciferase control (shLuc). Relative cell survival was normalized to untreated controls to give percent survival. Error bars indicate s.d. (E-F) MMC and CPT cell survival assay of isogenic BJ BRCA2^8488-1G>A^ or BRCA2^8524C>T^ clones depleted of either SMARCAL1 or ZRANB3 by shRNA or transduced with shRNA luciferase control (shLuc). Relative cell survival was normalized to untreated controls to give percent survival. Error bars indicate s.d.

### SMARCAL1, ZRANB3, and HLTF function is not required for ICL repair

Replication fork reversal has been observed as a response to replication stress induced by a number of different classes of genotoxic agents including MMC [39]. SMARCAL1, ZRANB3, and HLTF are ATPase dependent DNA translocases of the SNF2 family of chromatin remodelers that have recently been shown to promote replication fork reversal *in vitro* and *in vivo*. Depletion of any of the three translocases rescues nascent strand resection at HU stalled forks in *BRCA2* deficient cells [29, 40]. Similarly, depletion of the translocases in the *BRCA2^8524C>T^ and BRCA2^8488-1G>A^* mutants rescued nascent strand degradation **(Figure 5A).** However, depletion of SMARCAL1, ZRANB3, or HLTF did not rescue increased RPA activation and foci formation after MMC **(Figure 5B-C**, Figure S5G-I). To determine if the proteins implicated in replication fork reversal are important for the repair of ICLs, wild type cells were depleted of SMARCAL1 or ZRANB3 and tested for sensitization to MMC. Cells depleted of either translocase were not significantly sensitized to MMC **(Figure 5D).** Additionally, depletion of either translocase did not rescue cellular hypersensitivity to MMC or CPT in *BRCA2^8488-1G>A^* cells **(Figure 5E-F).** These data suggest that the function of these translocases is not required during ICL repair.

## Discussion

### BRCA2 and RAD51 function at the ICL

Here we have studied the functional consequences of pathogenic *BRCA2* variants in the DNA binding domain in the context of homologous recombination, and protection of stalled replication forks due to dNTP depletion or DNA interstrand crosslink lesions. The DBD variants did not affect IR sensitivity, SCE levels, or HU sensitivity suggesting that the HR in cells carrying the DBD variants is largely intact. We also saw only a moderate reduction in HR using an HR reporter assay. Similar to the previously described patient cell line with RAD51/FANCR p.T131P mutation [27], the cells with BRCA2 DBD variants were sensitive to ICL inducing agents and showed increased RPA foci formation after MMC that was DNA2-WRN dependent. These data suggest that like the well described interdependence of BRCA2 and RAD51 in HR, BRCA2 and RAD51 function together in the early steps of ICL repair to prevent DNA resection and that the function of the BRCA2 DBD is important for this role. This expands the role of BRCA2 in ICL repair beyond HR to include protection of DNA at the ICL-stalled replication fork from aberrant nucleolytic processing (Figure 6).

**Figure 6:**
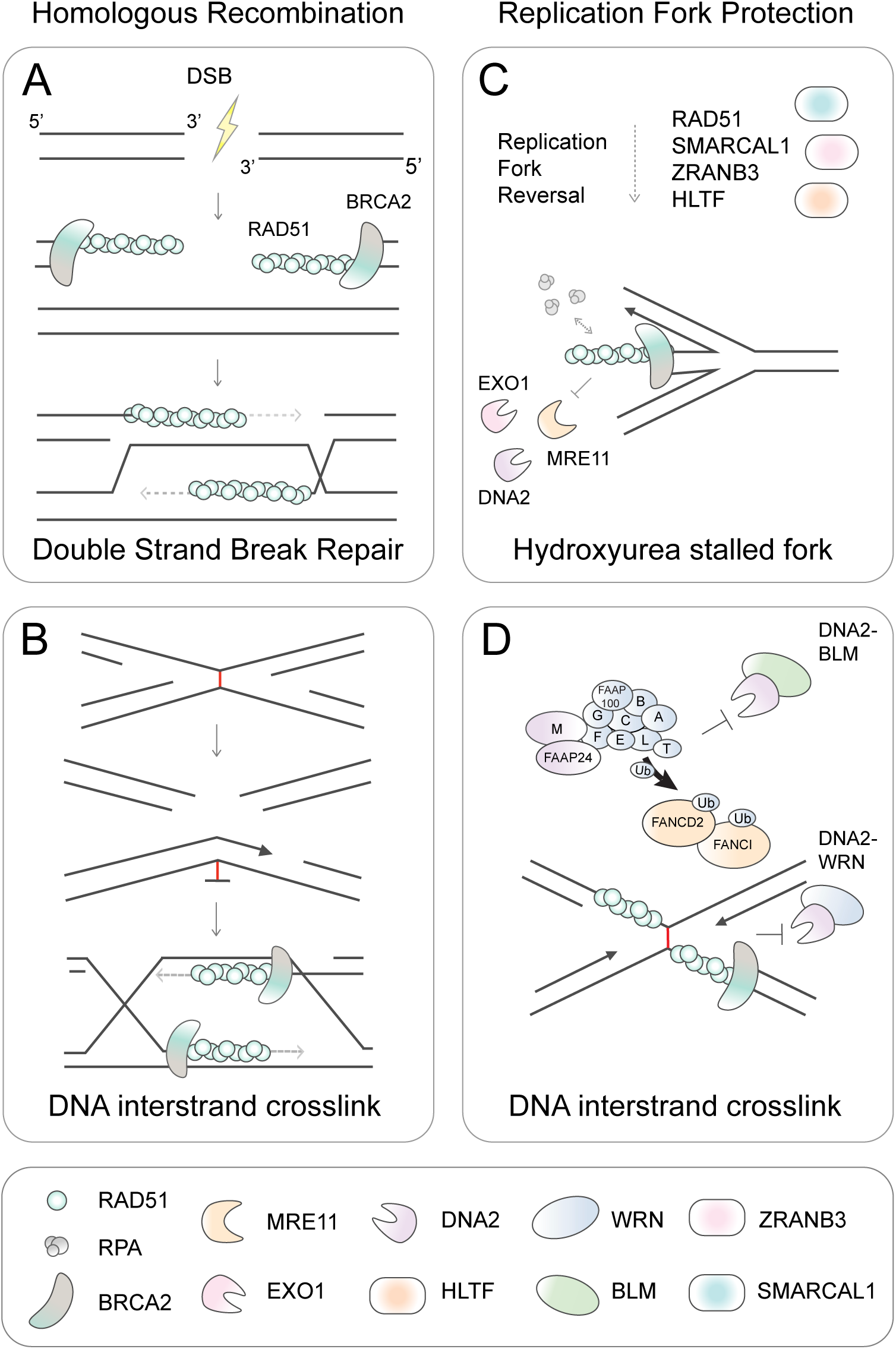
The role of BRCA2 in response to replication stress produced by hydroxyurea and DNA interstrand crosslinks is distinct. Schematic representing the different roles of BRCA2 in replication fork protection and homologous recombination. (A) During homologous recombination repair of DSBs, BRCA2 assembles RAD51 nucleofilaments onto ssDNA overhangs, which is important for the RAD51 mediated homology search of the sister chromatid. (B) During DNA interstrand crosslink repair homologous recombination is required to repair the programmed DSBs. BRCA2 has a role in two distinct types of replication fork protection. (C) At HU stalled forks, replication fork remodeling depends on RAD51 and the SNF2 translocases, SMARCAL1, ZRANB3, and HLTF. BRCA2 and RAD51 protect the reversed replication fork from degradation by nucleases. The MRE11 nuclease has been reported numerous times to be responsible for the degradation of HU-stalled forks in the absence of fork protection. More recently other nucleases have been described in nascent strand degradation including EXO1 and DNA2. (D) At ICLs, BRCA2 and RAD51 protect the fork from resection by the DNA2-WRN nuclease helicase complex. The ssDNA generated after MMC is not dependent upon MRE11, CTIP, or EXO1, as described for HU-stalled forks. ICL repair does not require the function of the SNF2 translocases suggesting that reversed forks present in MMC treated cells are likely the result of a more global cellular response to replication stress. We also propose that an additional role of the FA core complex and associated proteins at the ICL is to prevent aberrant resection by the DNA2-BLM nuclease helicase complex.

Depletion of the replication fork remodelers SMARCAL1, ZRANB3, and HLTF and the RAD51 modulator RADX rescued nascent strand degradation at HU-stalled forks in cells carrying DBD variants consistent with the previous data on the role of BRCA2 in this process [25, 29, 38, 40]. However, depletion of the translocases did not mitigate cellular sensitivity or increased RPA after MMC in the BRCA2 DBD mutant cells. Our study demonstrates that remodeling by the translocases is not a major step in the repair of ICLs and suggests that the MMC induced replication fork reversal may be a more general response to replication stress but not specifically at the fork that is stalled at an ICL [39, 41]. These data further support that the protection by BRCA2 and RAD51 at a HU-stalled fork is different from protection at an ICL (Figure 6).

The mechanism of the DNA protection at the ICL by the BRCA2 DBD domain remains to be explored. However, the location of the variants at the transition of the OB2 fold and base of the Tower domain suggests a plausible mechanism of protection at an ICL-stalled fork. OB2 fold binds to ssDNA and the Tower domain contains a 3HB domain at the apex that is capable of binding to dsDNA [42]. We speculate that the mutations in this region of the DBD may preclude efficient binding/bridging at ssDNA-dsDNA junctions which is a structure expected at stalled forks and lack of this binding would lead to the deprotection phenotype. Lack of proper placement of BRCA2 may also preclude proper RAD51 loading which also may lead to inappropriate DNA resection. Biochemical analysis of the BRCA2 variants we have identified in atypical Fanconi anemia patients will further our understanding of how BRCA2 interacts with different replication fork structures.

### FA protein function at the ICL

FA proteins have previously been shown to be important for protection at HU-stalled replication forks [43]. Here we show that FA patient cell lines from various complementation groups also demonstrate increased ssDNA and RPA foci formation after MMC. However, in FANCA deficient cells, the increase in RPA foci is dependent on DNA2 and BLM, but not WRN. This suggest that the fork protection of BRCA2-RAD51 is not redundant with the FA core complex, but further investigation will be needed to determine the genetic dependency of increased ssDNA in the absence of the other FA proteins. DNA2 has previously been reported to interact with FANCD2 and be recruited to ICLs where it is required for repair, but is deleterious in the absence of FANCD2 [44, 45]. BLM has been reported to interact with a number of FA proteins and co-localize with FANCD2 at ICLs [46–48]. Consistent with BLM depletion rescuing increased ssDNA at the fork in the absence of FANCA, BLM knockout was also recently reported to rescue ICL sensitivity and reduce DNA damage in FA deficient cells [49]. It is possible that DNA2, WRN, and BLM are recruited to ICLs for normal functions, but in the absence of key FA/BRCA pathway components are left unregulated resulting in aberrant processing of the fork.

### BRCA2 function at the HU-stalled replication fork

Our analysis of BRCA2 DBD mutants demonstrates that the function of the DBD is also required for protection at HU-stalled replication forks to prevent nuclease degradation. This is in contrast to a previous report that the DBD was dispensable for replication fork protection at HU-stalled forks [28]. Thus, the replication fork protection role of BRCA2 at HU-stalled replication forks is not restricted to the C-terminal domain and that fork protection likely requires the DBD to bind DNA at the stalled replication fork. It remains to be determined if the DBD variants have an effect on replication fork reversal after HU treatment but the dependency on the resection phenotype on the translocases suggest that they will.

While the role of MRE11 in nascent strand degradation of BRCA2 deficient cells has been widely shown, there is conflicting data about resection mediated by DNA2 [25, 26, 50]. A role for DNA2 with WRN in replication fork restart has been described, and it has also been reported that DNA2 degrades nascent DNA at stalled forks in the setting of RECQ1, BOD1L, or Abro1 deficiency [24, 51–53]. Here we show in isogenic cell lines that BRCA2 function is required to also prevent DNA2 resection at HU-stalled forks.

The observation that genomic instability results from the absence of proper replication fork protection after HU treatment has largely been studied by RNAi depletion of BRCA2 [25, 29, 40]. By studying *BRCA2* mutants, we show that a significant increase in chromosomal breakage after HU does not correlate with replication fork resection. For some of the BRCA2 mutants (c.8524C>T and p.S3291A) replication fork protection at HU-stalled forks is defective, but there is no significant increase in chromosomal breakage after HU. We observed increased chromosomal breakage in cells expressing BRCA2 LOF truncation variant, which is consistent with many previous reports that BRCA2 knockdown results in increased chromosomal breakage [25, 28, 29, 40]. The DNA damage in cells with BRCA2 LOF variants was similarly rescued by MRE11 depletion/inhibition. However, all of the *BRCA2* mutants in our analysis that undergo MRE11 dependent fork resection at HU-stalled replication forks do not have significantly elevated chromosomal breakage. These results also correlate with the cellular sensitivity observed in the BRCA2 mutants; LOF mutants show sensitivity to replication stress induced by HU and aphidicolin whereas the DBD mutants did not. These results demonstrate the importance of using *BRCA2* mutants that permit separation between different BRCA2 functions as opposed to RNAi depletion or LOF mutants that remove all function. It is possible that in these depletion studies or with LOF mutants, that the loss of BRCA2 HR function contributes to the breakage phenotype at the unprotected and degraded replication forks.

We show that DNA2 depletion in BRCA2 mutant cells also rescues resection at HU-stalled forks, but at the same time we observe that DNA2 depletion exacerbates chromosomal breakage after HU treatment. This observation suggests that in the setting of BRCA2 deficiency DNA2 depletion is deleterious, which may be due to a requirement in replication-coupled repair or modulation of reversed forks [44, 52, 54]. Recent reports have also implicated EXO1 and CTIP as degrading HU-stalled forks in the absence of BRCA2 [25]. Conversely, CTIP has been reported to be required to restrain DNA2 activity at stalled replication forks in the absence of BRCA1/2 [26]. Taken together, resection of the regressed fork in the absence of BRCA2 is now reported to involve all of the DSB end-resection nucleases. MRE11 and DNA2 are already reported to be required for replication fork restart [52, 55]. However, further investigation is required to determine if all of these factors have a normal function in processing stalled forks or restoring reversed forks under wild type genetic conditions. These results are also interesting in that all of the nucleases are implicated at HU-stalled forks, but only DNA2 has activity at the ICL in BRCA2 deficient cells.

### Clinical implications

The identification of BRCA2 DBD variants in conjunction with atypical disease presentation gives the opportunity to investigate how defects in the DBD impact BRCA2 function and gives insight into how these defects may give rise to the developmental defects characteristic of FA but not the early childhood malignancies seen in other patients with biallelic *FANCD1/BRCA2* variants. The disease presentation of these individuals resembles the phenotype of FA-like patients described for FA-R (FANCR/RAD51) and FA-O (FANCO/RAD51C) complementation groups [22, 27]. Due to the moderate impact that these DBD variants have on HR, we hypothesize that the retention of ∼50% of HR function that we observe is sufficient enough to safeguarded against early tumor development. Diagnosis and classification as FA-D1 (BRCA2/FANCD1) complementation group should also be considered for patients presenting with FA-like syndrome.

Furthermore, this study has implications for how we think about *BRCA2* variants of unknown significance (VUS) in human disease, including HBOC and FA. Evaluation of BRCA2 VUS relies on multifactorial probability models [56] or functional assays assessing HR [57] to estimate if a variant is pathogenic. Some VUS can be easily classified as pathogenic if HR is dramatically reduced; however, a number of VUS show intermediate phenotypes, making it difficult to interpret their role in HBOC [57, 58]. The contribution of other BRCA2 functions, including replication fork protection, to cellular function and tumorigenesis requires further investigation. We have demonstrated that these BRCA2 DBD mutations are pathogenic. It is possible that some mutations carry only low to moderate risk for HBOC related to preservation of HR function, but still result in FA when biallelic BRCA2 mutations are inherited due to a predominant defect in ICL repair.

## Acknowledgements

We thank the family and patients for their participation in the IFAR and this study. We thank Alberto Ciccia for sharing shRNAs to SMARCAL1 and ZRANB3. We thank Alan D’Andrea for sharing HSC62 cells. We thank the A.S. laboratory members for their input and discussion of the manuscript. Initial BRCA2 studies were funded by a Starr Foundation grant I8-A8-101. RAD51 and BRCA2 studies in the laboratory are supported by grant RO1 CA204127 from NIH and grant # UL1TR001866 from the National Center for Advancing Translational Sciences, NIH Clinical and Translational Science Award program. KAR was supported by the Medical Scientist Training Program grant from the National Institute of General Medical Sciences of the NIH under award number T32GM007739 to the Weill Cornell/Rockefeller/Sloan-Kettering Tri-Institutional MD-PhD Program and the William Randolph Hearst Foundation Fellowship at the Rockefeller University. A.S. is an HHMI Faculty Scholar.

## Author Contributions

KAR and AS conceived the ideas and designed experiments for this study. KAR, RN, FPL, SS, ATW performed the experiments. AA and KAR performed bioinformatic analysis of the exomes. ADA directed clinical testing in the proband at the time of IFAR enrollment and excluded number of complementation groups in the patient. MK provided clinical information. KAR and AS wrote the manuscript with essential input from other authors. AS acquired funding and supervised all studies.

## Experimental Procedures

### Study subjects

DNA samples and cell lines were derived from subjects enrolled in the International Fanconi Anemia Registry (IFAR) after obtaining informed written consent. The Institutional Review Board of The Rockefeller University, New York, NY, USA, approved these studies.

### Cell lines

Patient-derived fibroblast cell lines (**Table S1**) and BJ foreskin normal control fibroblasts (ATCC) were transformed by expression of HPV16 E6E7 and immortalized with the catalytic subunit of human telomerase (hTERT). Fibroblasts were cultured in Dulbecco Modified Eagle medium (DMEM) supplemented with 15% FBS, 100 units of penicillin per mL, 0.1 mg of streptomycin per mL, non-essential amino acids, and glutamax (Invitrogen). Fibroblasts cell lines were incubated at 37°C, 5% CO_2_, and 3% O_2_. Lymphoblast cell lines (**Table S1**) were established from patient peripheral blood mononuclear cells by Epstein-Barr Virus (EBV) transformation and grown in Roswell Park Memorial Institute medium (RPMI) with 20% FBS and further supplemented as above. HEK293T (ATCC) cells were cultured in DMEM supplemented with 10% FBS and penicillin/streptomycin and glutamax as indicated above. Lymphoblast and HEK293T cell lines were incubated at 37°C, 5% CO_2_, and ambient O_2_.

### Viral transfection/transduction

cDNAs were delivered by retroviral or lentiviral transduction after packaging in HEK293T cells (TransIT-293 transfection reagent, Mirus). HEK293T cells were plated at 4.5*10^6^ the evening before transfection of DNA and viral packaging vectors. Transfection was performed according to the manufacturer’s instructions. The next day after transfection cell media was replaced and two days after transfection viral supernatants were harvested and used to infect target cells in the presence of 4 mg/ml polybrene. Stably expressing cells were selected with the appropriate agent puromycin (2 μg/ml), hygromycin (100-200 μg/ml), blasticidin (500 μg/ml), neomycin (600 μg/ml).

### RNAi

Cells were transfected with pools of 3 siRNAs against MRE11, DNA2, EXO1, CTIP, WRN, BLM, BRCA2, RAD51, MUS81, XPF, and SLX4. For RADX and HLTF depletion a single previously published siRNA was used (**Table S2**) [38, 40]. Cells were transfected using Lipofectamine RNAiMAX (Invitorgen) according to the manufacturer’s instructions. For shRNA depletion, virus was packaged in HEK293T cells and used to infect target cells and cells with stable integration were selected. shRNA constructs for SMARCAL1 and ZRANB3 were a gift from Alberto Ciccia (**Table S3**). shRNAs to *RADX* were purchased from Transomics and used in the pZIP_hCMV_Puro vector or pMSCV-PM-mir30. shRNAs were PCR amplified and cloned into pMSCV-PM-mir30 by digestion with XhoI and MluI and vector ligation. See **Table S4** for PCR primers for amplification of shRNA from UltramiRs of pZIP_hCMV vector. RNAi knockdown was measured by RT qPCR or western blot.

### PCR, reverse transcription, and RT qPCR

PCR reactions were performed using Taq DNA Polymerase (Qiagen), Phusion High-Fidelity PCR Master Mix with GC buffer (Thermo Scientific), and PCR SuperMix High Fidelity (Invitrogen) according to manufacturer’s protocols and primers are listed in **Table S5**. Total messenger RNA was extracted using RNeasy plus kit (Qiagen). RNA was reverse transcribed to cDNA using the SuperScript III Reverse Transcriptase (Invitrogen). Platinum SYBR Green SuperMix-UDG (Invitrogen) was used according to manufacturer’s protocol to determine relative transcript levels which were normalized against GAPDH levels. (See **Table S6** for RT qPCR primers).

### Gene targeting

To correct the *BRCA2* c.8488-1G>A variants in HSC62 fibroblasts, cells were transduced with the pCW-Cas9-Puro (addgene #50661) vector which contains a doxycycline inducible Cas9. Subsequently, HSC62 cells were transduced with plentiGuide-Hygro (derived from addgene #52963) that expresses a single guide RNA (sgRNA) (**see Table S7** for sgRNA sequence) that targets DNA in proximity to the c.8488-1G>A variant. sgRNAs were designed using the online CRISPR design tool from the Zhang laboratory (crispr.mit.edu). 1*10^6^ cells were electroporated with a 100bp template oligonucleotide (see **Table S8** for sequence) using Lonza 2b-Nucleofector. Cells were cultured in 500 ng/mL doxycycline for 48 hours to induce Cas9 expression and then incubated in fresh doxycycline free media for another 48 hours before being single cell cloned into 96-well plates. Clones were expanded and screened by sequencing of genomic DNA. For clones HSC62^mut/WT^-1and HSC62^WT/WT^-2, cells were selected in low dose MMC (50 ng/mL) once a week for three weeks before seeding in 96-wells. Clone 3 (HSC62^WT/WT^) was not selected for.

The rest of the gene targeting was performed by electroporation of Cas9/gRNA ribonucleoprotein (RNP) complexes with 100nt oligonucleotide donor templates, with phosphorothioate protected ends. sgRNA was prepared by combining crRNA (designed using crispr.mit.edu) and universal tracrRNA as per manufactures guidelines (IDT). To form RNP complexes gRNA duplex and Cas9-3NLS (IDT) were combined, incubated at room temperature for 10-15 minutes, and then placed on ice until used. RNP complexes and 10 ug of 100nt donor template oligonucleotide were electroporated into 2*10^5^ fibroblasts or 3.5*10^5^ HEK293T cells using Lonza 4D-Nucleofector. Cells were plated in a 12-well for 48-72 hours to recover before single-cell plating in 96-wells. Clones were expanded and screened by sequencing of genomic DNA. No selection was used.

### Chromosomal breakage

Cells were treated with 0.1 μg DEB per mL of media for 48-72 hours or 45-100 nM of MMC for 24 hours. HU and aphidicolin treatments were as indicated. LCLs were arrested with colcemid (0.17 μg/mL) for 20 minutes and fibroblasts for 90 minutes. Cells were harvested and incubated in 0.075 M KCL for 10 minutes before being fixed in methanol and acetic acid (3:1). Cells were dropped onto wet slides and dried at 40°C for at least one hour before staining with Karyomax Giemsa (Invitrogen) for three minutes. Dry slides were then imaged on the Metasystems Metafer slide scanning platform.

### Cell survival studies

Fibroblasts were seeded overnight in triplicate and treated the next day with DNA damaging agents at indicated concentrations. Cells were grown for 4-6 days and passaged once at appropriate ratios. Once cells reached near confluence (7-9 days), cells were counted using Z2 Coulter counter (Beckman Coulter). In the case of cisplatin treatment, drug was removed after 1 hour and cells were washed with PBS and given fresh drug-free media. For aphidicolin treatment, after 48 hours cells were washed with PBS and given fresh drug-free media. For PARPi treatment, cells were given fresh media with olaparib daily. For ionizing radiation cells were treated with the indicated IR dose in Falcon tubes prior to being plated. LCLs were treated at the time of seeding, agitated daily, and counted on the 7^th^ day. HEK293T cells were seeded overnight, treated with MMC, passaged after 3 days, and counted on the 5^th^ day.

### Western blot

Whole cell extracts were prepared by lysing cell pellets in Laemmli sample buffer (Bio-Rad or 4% SDS, 20% glycerol, 125 mM Tris-HCl pH 6.8). Samples were either sonicated or vortexed at highest speed for 30 seconds. Samples were boiled for 5 minutes. For pRPA and BRCA2 western blots, samples were instead heated at 50°C for 10 minutes. Proteins were separated on 4-12% or 3-8% gradient gels (Invitrogen) by SDS-PAGE. Immunoblotting was performed using the antibodies indicated in **Table S9**.

### Immunofluorescence

Cells were seeded on coverslips the day before. For FAND2 foci, cells were treated with 1 μM MMC for 24 hours. For RAD51 foci, cells were irradiated for indicated dose or treated with 3 μM MMC for 1 hour and harvested at indicated times. For RPA foci cells were treated with 3 μM MMC for 1 hour and harvested at indicated times. Cells were washed with PBS twice, fixed in 3.7% formaldehyde for 10 minutes, washed twice with PBS, and permeablized with 0.5% Triton in PBS for 10 mins. Cells were blocked in 5% [v/v] FBS in PBS and incubated with primary antibodies in blocking buffer for two hours at room temperature or overnight at 4°C (for antibodies see **Table S9**). Cells were washed three times for five minutes with blocking buffer and then incubated with secondary antibody (1:1000) (Alexa Fluor). Cells were washed again three times with blocking buffer, rinsed quickly with water, air dried, and then embedded on glass slides with DAPI Fluoromount-G (SouthernBiotech).

### Sister chromatid exchange

For MMC induced SCEs, fibroblasts were cultured for 24 hours in 10 ug/mL BrdU and then treated with 0.1 or 0.2 ug/mL MMC for one hour. Cells were washed and put into fresh media with 10 ug/mL BrdU for another 24 hours. For cells depleted of BLM, siRNA transfection was performed twice as described. For the second siRNA transfection 10 ug/mL BrdU was added to media and cells were cultured in BrdU for a total of 48 hours before harvest. Cells were collected, fixed, and dropped on glass slides for metaphases as previously described. Slides were dried overnight at 42°C and then stained in 20 ug/mL Hoechst 33342 for 30 minutes. Slides were treated with 254 nM UV light for 3 hours. Slides were incubated at 65°C in 2x SCC for 2 hours, then rinsed in 1x GURR buffer, and stained in 8% Giemsa Karyomax for 3 minutes. Metaphases were scanned and imaged on Metasystems Metafer Slide Scanning Platform.

### mClover homologous recombination assay

Cells were plated in a 24-well plate the day before and transfected with 0.25 ug pCMV-Cas9-sgLMNA-BFP and 0.4 ug pDONR-LMNA using TransIT-293 Transfection Reagent (Mirus) according to manufactures instructions (plasmids were a gift from Jan Karlseder) [34]. 24 hours after transfection cell media was replaced. Cells were incubated for another 48 hours and were then harvested and analyzed on BD LSRII to determine the proportion of mClover positive cells and data was analyzed with FlowJo.

### DNA fibers

For DNA fibers, cells were plated the evening before and labeled with nucleotide analogs and treated with 6 mM HU for five hours. Cells were harvested and cell pellets were washed one time in cold PBS. Cells were resuspended at a concentration of 1*10^6^ cells/mL in cold PBS. On a clean glass coverslip 10 ul droplets of spreading buffer (0.5% SDS, 200mM Tris-HCl pH 7.4, and 50 mM EDTA pH 8) was placed. 2.5 ul of cell suspension was pipetted into the spreading buffer, stirred, and pipetted up and down three times. Coverslips were incubated horizontally for nine minutes at room temperature before gently being tilted vertically to allow the buffer to run down the slide. Coverslips were dried at room temperature at an angle and then heated at 65°C for 30 minutes. Coverslips were fixed in methanol/acetic acid 3:1 overnight at 4°C. The next day coverslips were washed in PBS three times at room temperature and then incubated in 2.5M HCl for 1 hour. Coverslips were then washed five times for five minutes with PBS and after the final wash they were blocked in 5% FBS in PBS for 30 minutes. For immunostaining, coverslips were incubated with primary antibodies for 2.5 hours at room temperature. Rat anti-BrdU antibody (1:40) was used to detect CldU and mouse anti-BrdU antibody (1:20) was used to detect ldU. Coverslips were washed 5 times with PBS with 0.2% Tween and then blocked for 30 minutes in 5% FBS in PBS. Coverslips were incubated with secondary (Alexa Fluor) anti-rat (594) and anti-mouse (488) at a dilution of 1:300 for 1 hour at room temperature. Coverslips were washed 5 times with with PBS with 0.2% Tween and rinsed with water and air dried. Dry coverslips were mounted on glass slides using Fluoromount-G (SouthernBiotech). DNA tracks were all imaged on the DeltaVision Image Restoration microscope and measured using ImageJ.

### Whole Exome Sequencing

The libraries for whole exome sequencing (WES) were constructed and sequenced on Illumina HiSeq 2000 or Illumina GA-IIX using 76 bp paired-end reads at the Broad Institute or by using Agilent SureSelect Human All Exon V4 capture kit and 100 bp paired-end sequencing on Illumina HiSeq 2500. Sequence was aligned to human genome build GRCh37 using BWA (Burrows-Wheeler Aligner) [59]. Duplicate reads were marked using Picard [http://picard.sourceforge.net]. Genome Analysis Toolkit (GATK) was used for base quality score recalibration (BQSR), and local realignment around indels [60]. Variant discovery was performed in part by variant calling with GATK HaplotypeCaller and then joint genotyping with GATK GenotypeGVCFs. The variant call sets were then refined with Variant Quality Score Recalibration (VQSR) and VQSR scores helped discriminate low quality variants. Variant annotation was performed using SnpEff, VCFtools, and in-house software (NYGC) [61, 62]. All WES was analyzed with the NYGC sequence analysis pipeline.

**NCBI References**: BRCA2/FANCD1 RefSeq: NM_000059.3, Protein: NP_000050.2

## Supplemental information

### Supplemental Tables

**Table S1:** Patient Phenotypes

|  |  |
| --- | --- |
| **Family 1:** |  |
| Maternal allele: c.2330dupA<br>Paternal allele: c.8524C>T |  |
| RA3105<br>23 years old | microcephaly, craniofacial abnormalities, radial ray defect, thumb abnormalities, gastrointestinal malformation, ear abnormalities, palpebral fissures, high myopia |
| RA3106<br>20 years old | Craniofacial abnormalities, absent and hypoplastic thumb, gastrointestinal malformations, dysplastic sacrum, cardiac defects, horseshoe kidney |
| Family History | Breast cancer, skin cancer, colorectal cancer |
| Clinical breakage studies |  |
| Hematological status | No bone marrow failure or malignancy |
| **Family 2:** |  |
| Maternal allele: c.8488-1G>A (C.8488-1G>A)<br>Paternal allele: c.8488-1G>A (C.8488-1G>A) |  |
| HSC62 [16]<br>Deceased due to accident<br>unrelated to disease | Thumb abnormality<br>No bone marrow failure or cancer reported (status to the age of 30) |

**Table S2:**
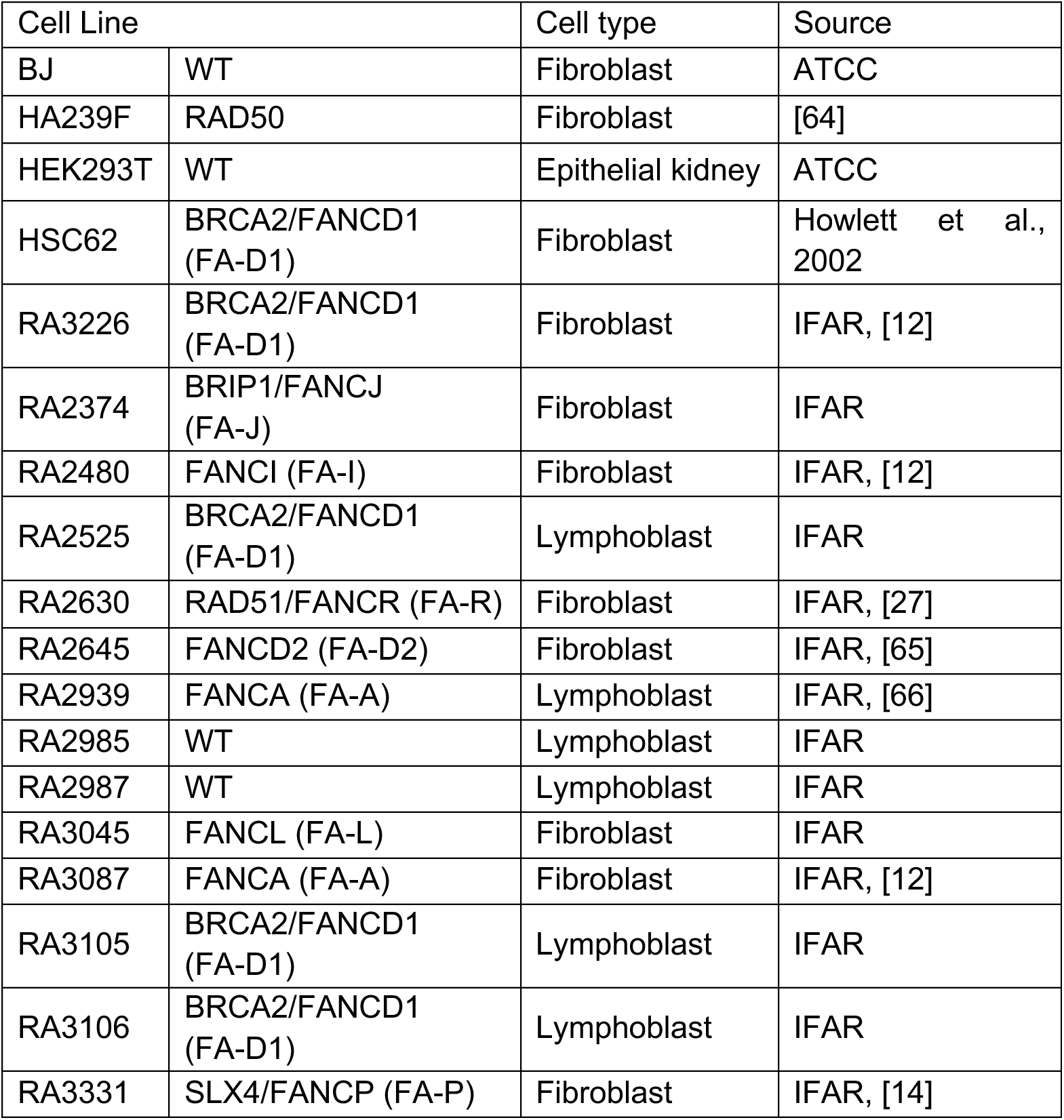
List of cell lines

**Table S3:**
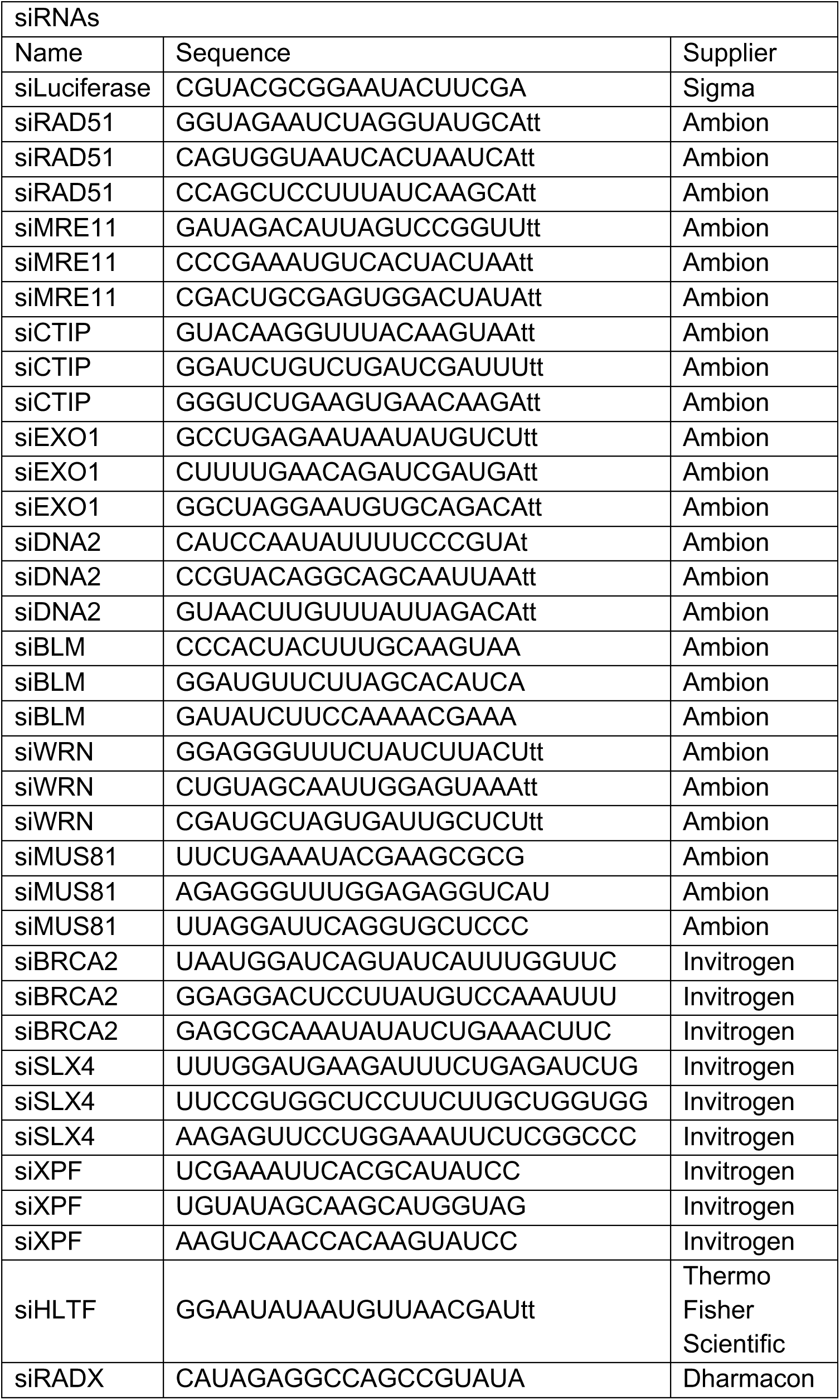
List of siRNAs

**Table S4:**
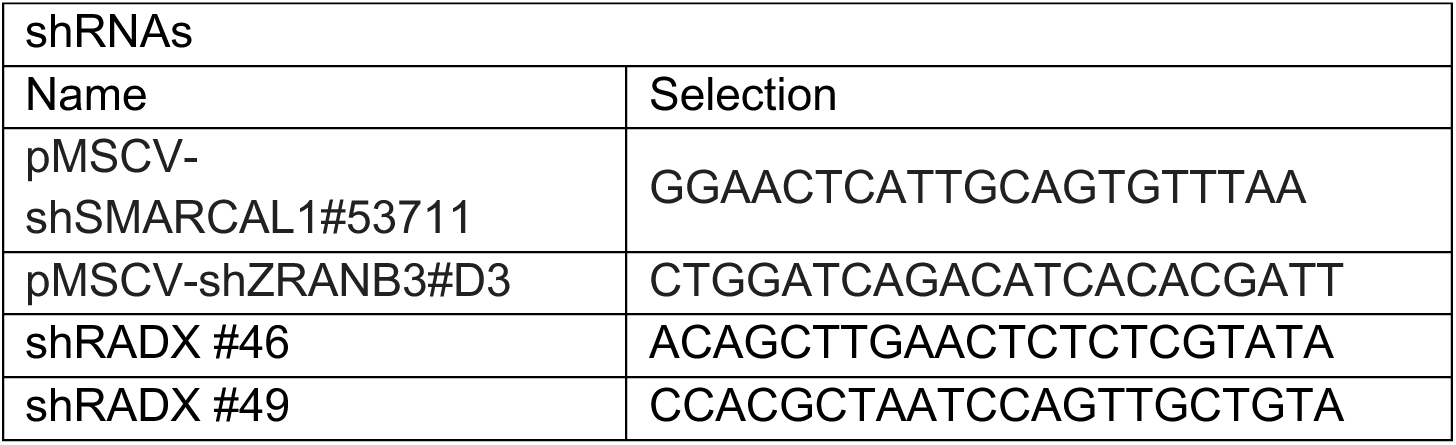
List of shRNAs

**Table S5:**
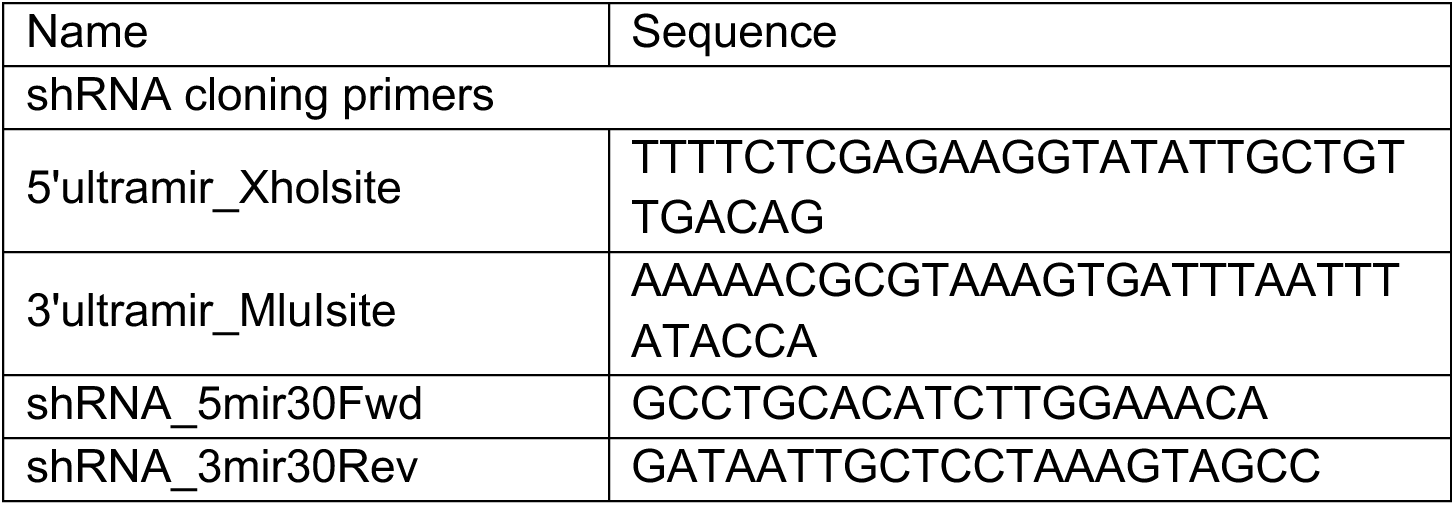
List of cloning primers

**Table S6:**
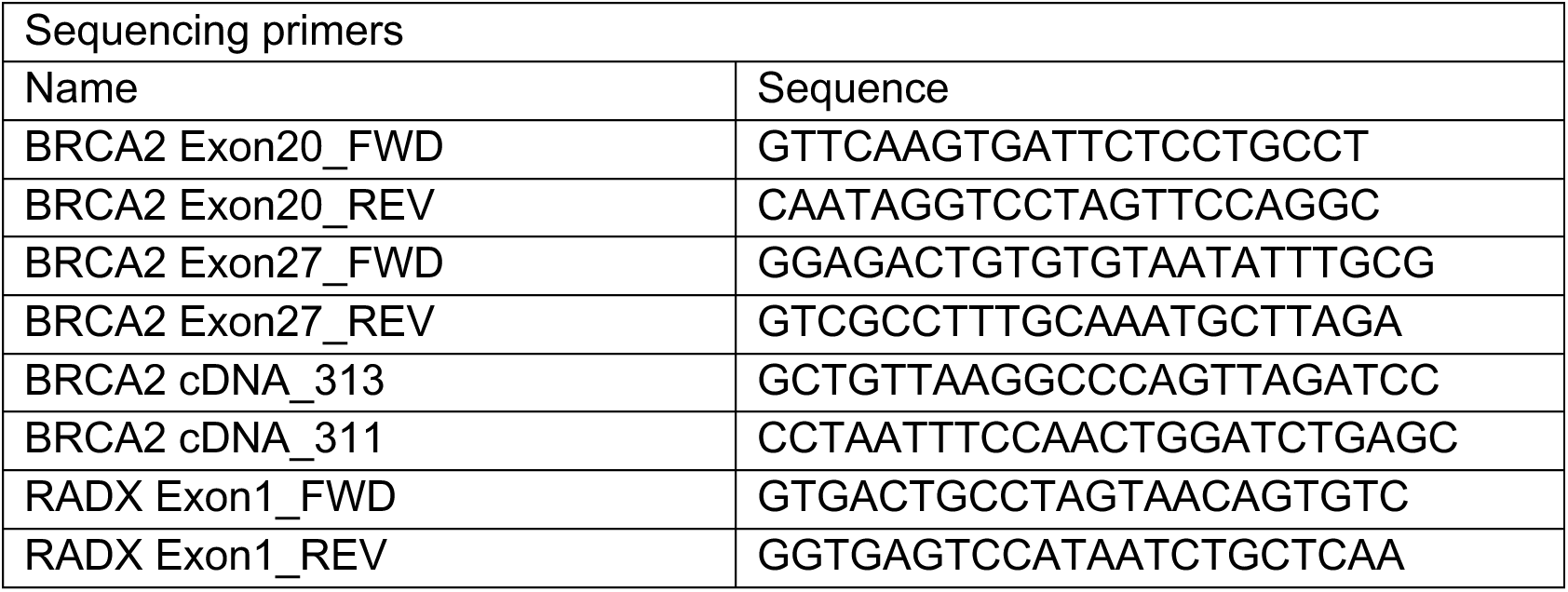
List of sequencing primers

**Table S7:**
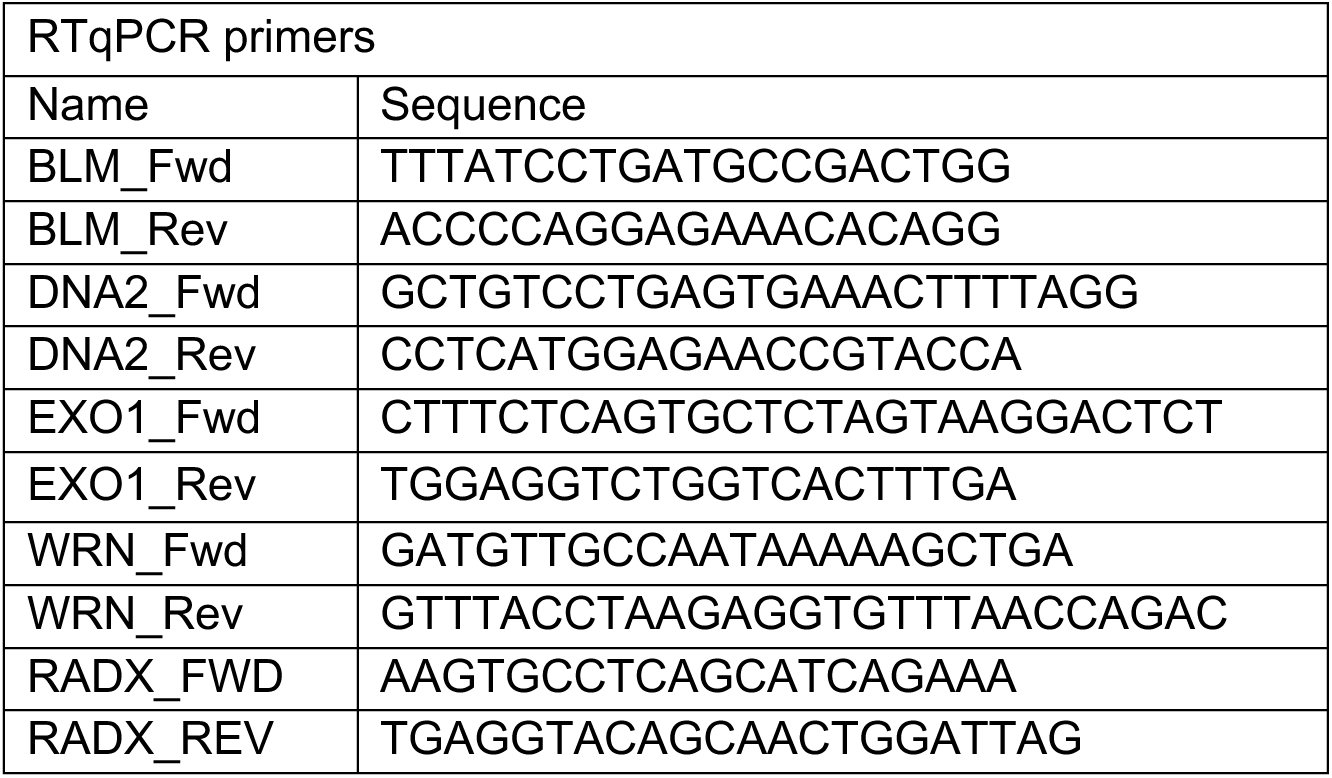
List of RT qPCR primers

**Table S8:**
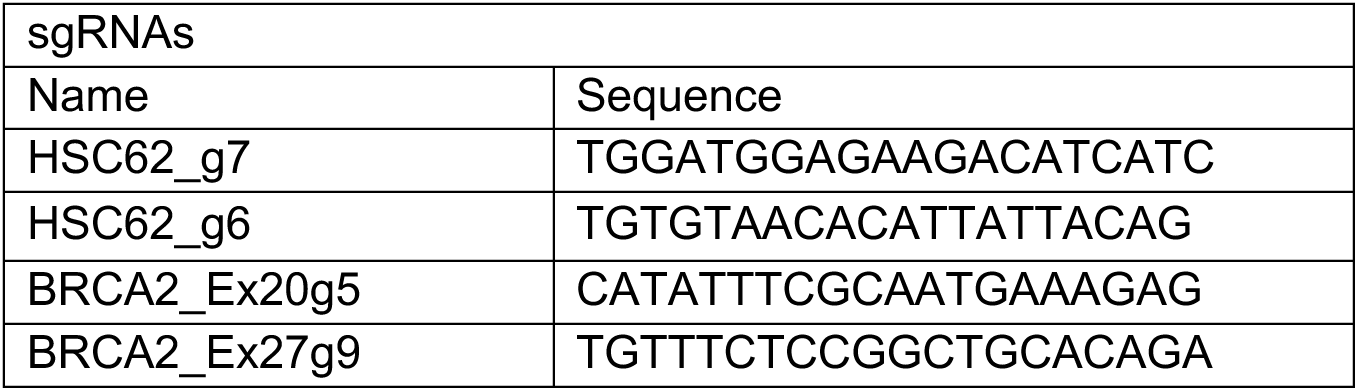
List of sgRNAs

**Table S9:**
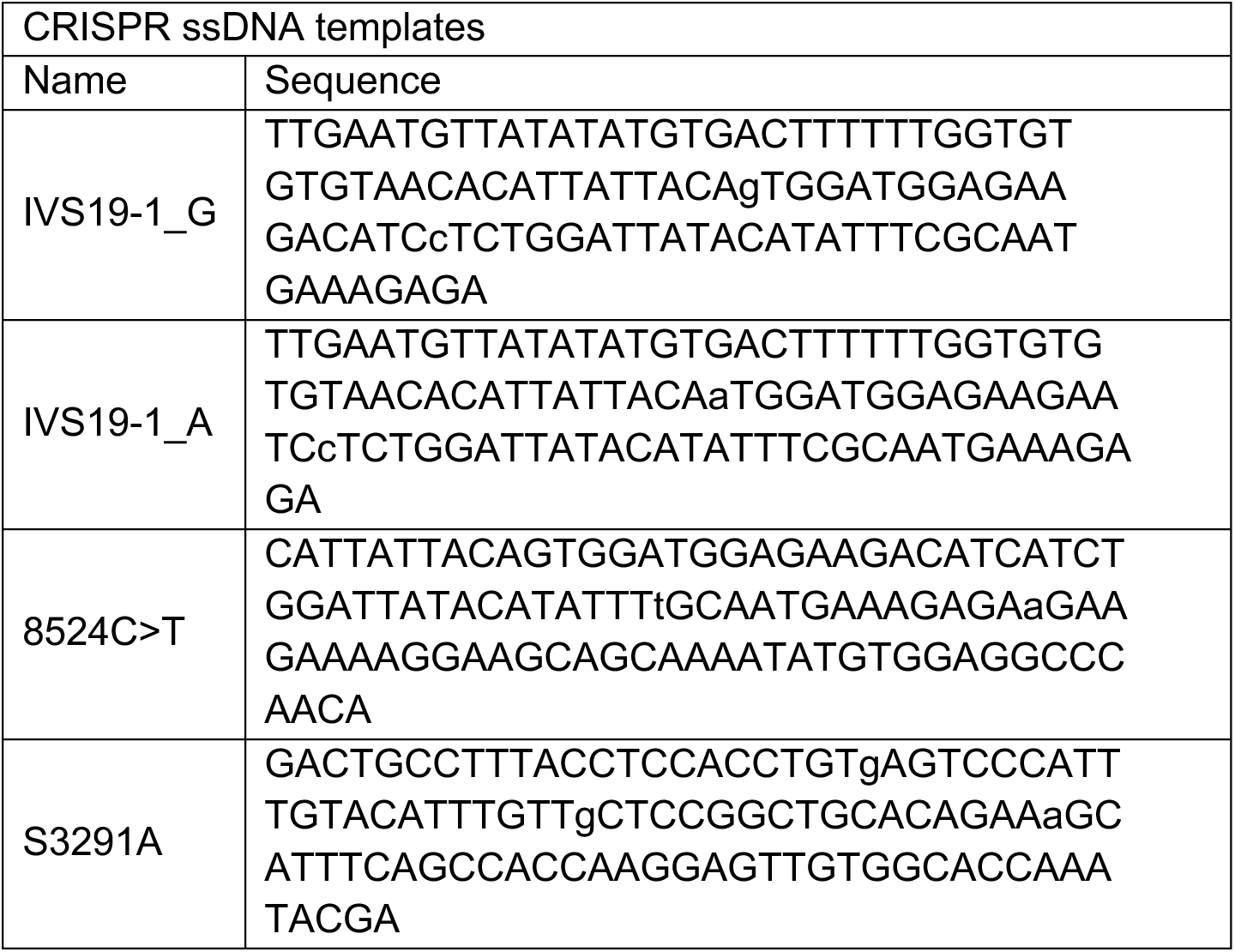
List of oligonucleotide donor templates for CRISPR/Cas9

**Table S10:**
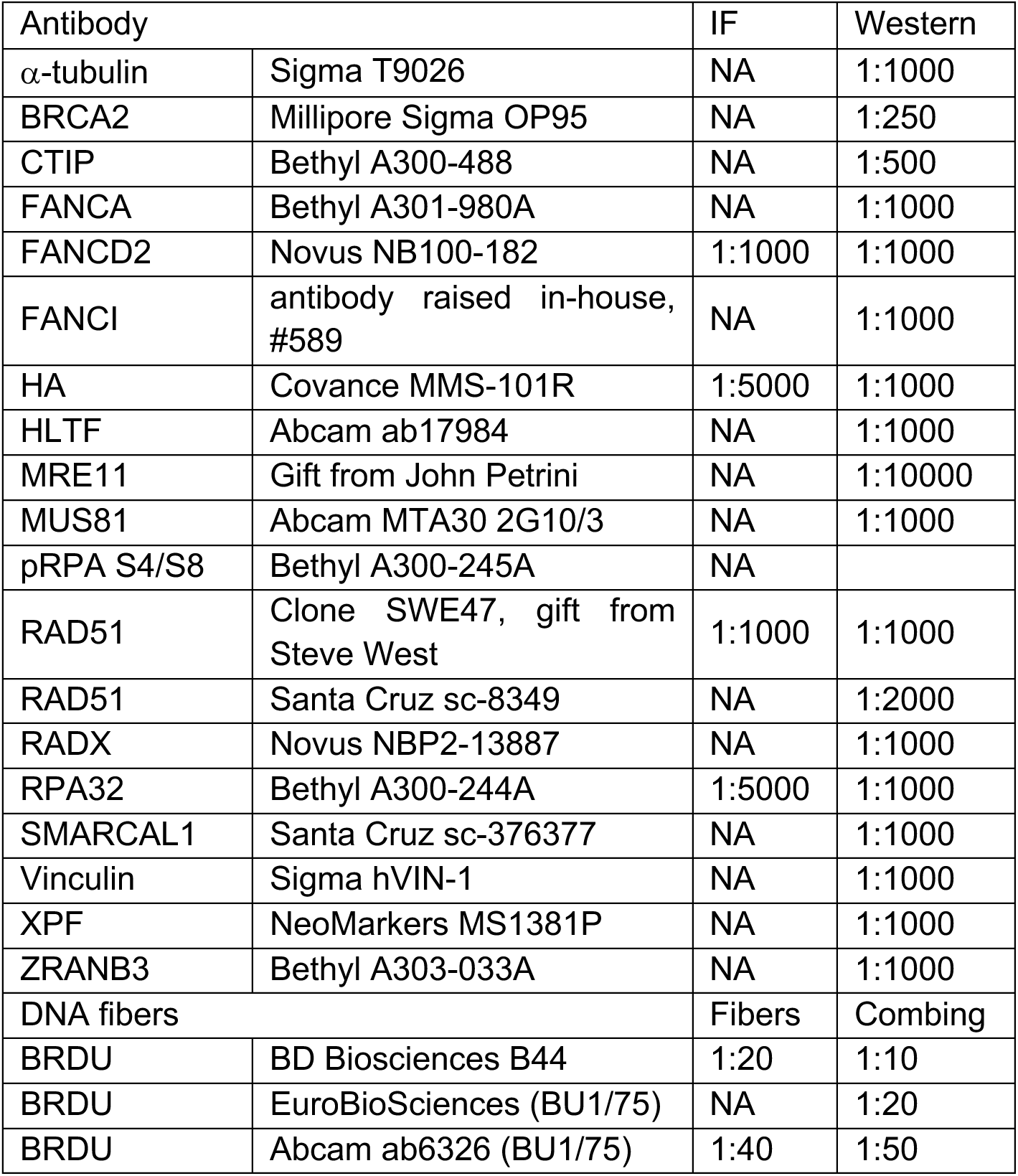
List of antibodies

**Figure S1:**
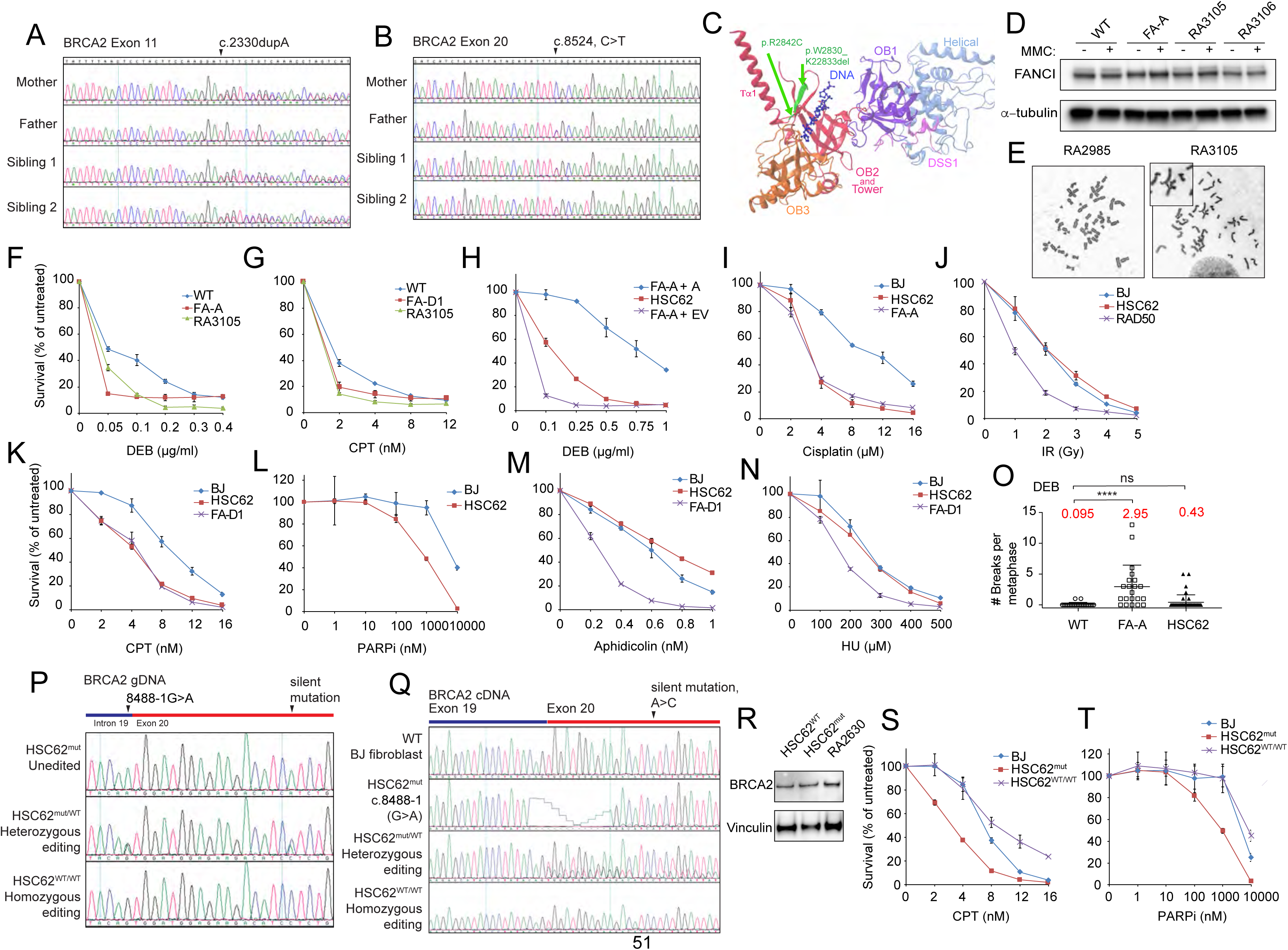
(A-B) Chromatograms of Sanger sequencing of DNA derived from a sibling pair and parents confirming the presence of *BRCA2* c.2330dupA and c.8524C>T variants identified by whole exome sequencing (WES). (C) BRCA2 structure of the DBD illustrating the location of p.W2830_K2833del and p.R2842C patient variants at the base of the Tower domain and OB2. Structure adapted from Yang *et al*., 2005. (D) Immunoblot analysis for FANCI ubiqutination following treatment with 1μM MMC for 24h of. WT (RA2985), FA-A (RA2939), and patient RA3105 and RA3106 LCLs. (E) Metaphase for RA2985 and RA3105 following DEB treatment. (F-G) Cell survival assays of patient derived lymphoblast cell line (LCLs) RA3105, FA-A (RA2939), WT (RA2985), and FA-D1 (RA2525) after (DEB) and camptothecin (CPT) treatment. Survival assays were performed in triplicate. Cells were treated with increasing concentrations of genotoxic agents and counted after 7-10 days in culture. Relative cell survival was normalized to untreated controls to give percent survival. Error bars indicate s.d. (H-N) Cell survival of HSC62 fibroblasts (c.8488-1G>A) to indicated agent compared to BJ WT fibroblast, FA-A patient fibroblast (RA3087), FA-A complemented patient cells expressing wild type *FANCA* (FA-A+A) or empty vector (FA-A+EV), RAD50 patient fibroblast or FA-D2 (BRCA2/FANCD1) patient fibroblast (FA-D1). Cell survival assays were performed in triplicate. Cells were treated with increasing concentrations of indicated agent. Cell survival was determined by counting cells after 7-9 days in culture. Relative cell survival was normalized to untreated controls to give the percent survival. Error bars indicate s.d. (O) Quantification of chromosome breaks following DEB treatment of BJ wild type fibroblasts, FA-A (FANCA) patient fibroblasts (FA-A^mut^), and HSC62 fibroblasts. (P) Chromatograms of PCR amplified gDNA of CRISPR/Cas9 targeted HSC62 fibroblasts. Gene editing reverted the c.8488-1G>A variant either to homozygous WT (HSC62^WT^) or heterozygous WT (HSC62^mut/WT^) at the endogenous locus in HSC62 patient cells. The silent variant that was incorporated to destroy the CRISPR PAM sequence is indicated. (Q)cDNA analysis of HSC62 clones with either homozygous or heterozygous correction of the c.8488-1G>A variant demonstrating rescue of the 12bp deletion of exon 20 that results from alternate splicing. (R) Immunoblot showing BRCA2 levels in CRISPR/CAS9 corrected patient cell line HSC62^WT^, uncorrected HSC62 cells (HSC62^mut^), and RA2630 FA-R (RAD51/FANCR) patient fibroblasts. (S-T) Cell survival of HSC62 uncorrected patient cell line (HSC62^mut^) compared to BJ WT fibroblast and CRISPR/Cas9 corrected wild type HSC62 (HSC62^WT^) clone. Error bars indicate s.d.

**Figure S2:**
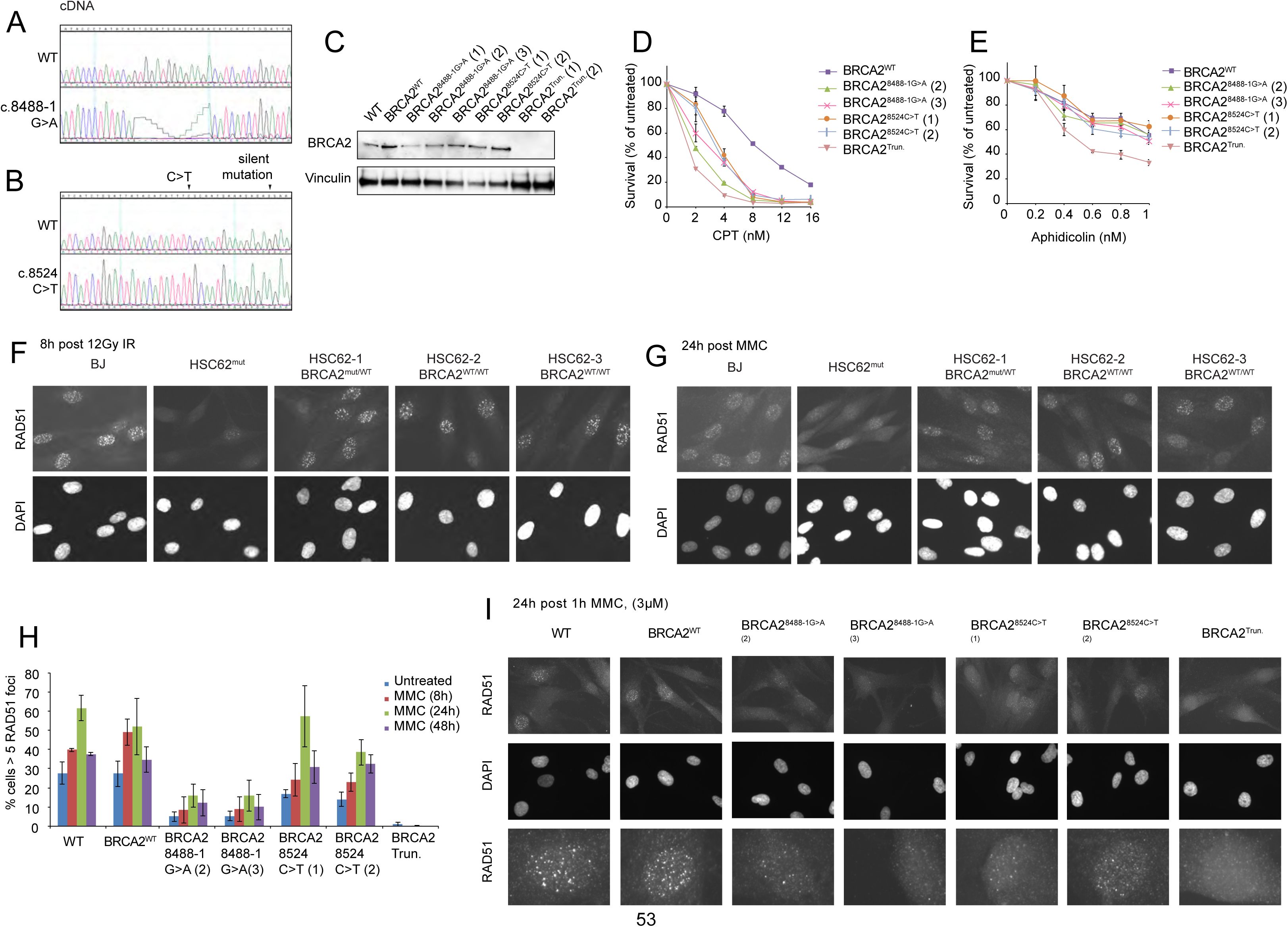
(A) cDNA sequencing of BRCA2 DBD CRISPR/Cas9 targeted BJ fibroblasts demonstrating that homozygous c.8488-1G>A variant in BJ fibroblast clones result in the usage of an alternative splice site donor and a 12bp deletion at the start of *BRCA2* exon 20 (as seen in patient derived HSC62 fibroblasts). (B) cDNA sequencing of BRCA2 DBD CRISPR/Cas9 targeted BJ fibroblasts demonstrating the c.8524C>T missense variant and silent mutation introduced by CRISPR/Cas9 targeting. (C) Immunoblot showing BRCA2 levels in bulk BJ WT fibroblasts, BRCA2^WT^ fibroblast clone, c.8488-1G>A BJ clones 1-3, c.8524C>T BJ clones 1-2, and *BRCA2^Trun.^* clones 1-2. (D-E) Cell survival of BJ WT fibroblasts, BJ WT fibroblast clone (BRCA2^WT^), c.8488-1G>A BJ clones, c.8524C>T BJ clones, and exon 20 *BRCA2* frameshift mutant (BRCA2^Trun.^). Cell survival assays were performed in triplicate. Cells were treated with increasing concentrations of CPT or aphidicolin. Cell survival was determined by counting cells after 7-9 days in culture. Relative cell survival was normalized to untreated controls to give the percent survival. Error bars indicate s.d. (F-G) Representative images of RAD51 foci in HSC62 uncorrected patient cell line (HSC62^mut^) compared to BJ WT fibroblast and CRISPR/Cas9 corrected wild type HSC62 (HSC62^mut/WT^ or HSC62^WT^) clones, 8h post IR and 24h post MMC. Detected by immunofluorescence with anti-RAD51antibody. Quantification in Figure 2C and 2D. (H) Quantification of RAD51 foci in isogenic BJ fibroblasts clones 8h, 24h and 48h following 1h treatment with 3 μM MMC of BJ WT fibroblasts, BJ WT fibroblast clone (BRCA2^WT^), c.8488-1G>A BJ clones 2-3, c.8524C>T BJ clones 1-2, and a BRCA2 truncation mutant, c.8531dupA (*BRCA2^Trun^*). Error bars indicate s.d. of three independent experiments (≥200 cells per experiment). (I) Representative images of RAD51 foci in isogenic BJ fibroblasts clones, 24h post 1h treatment with 3 μM MMC, detected by immunofluorescence with anti-RAD51antibody. Third row images are individual cells enlarged to better demonstrate differences in RAD51 foci size. Quantified in Figure S2H.

**Figure S3:**
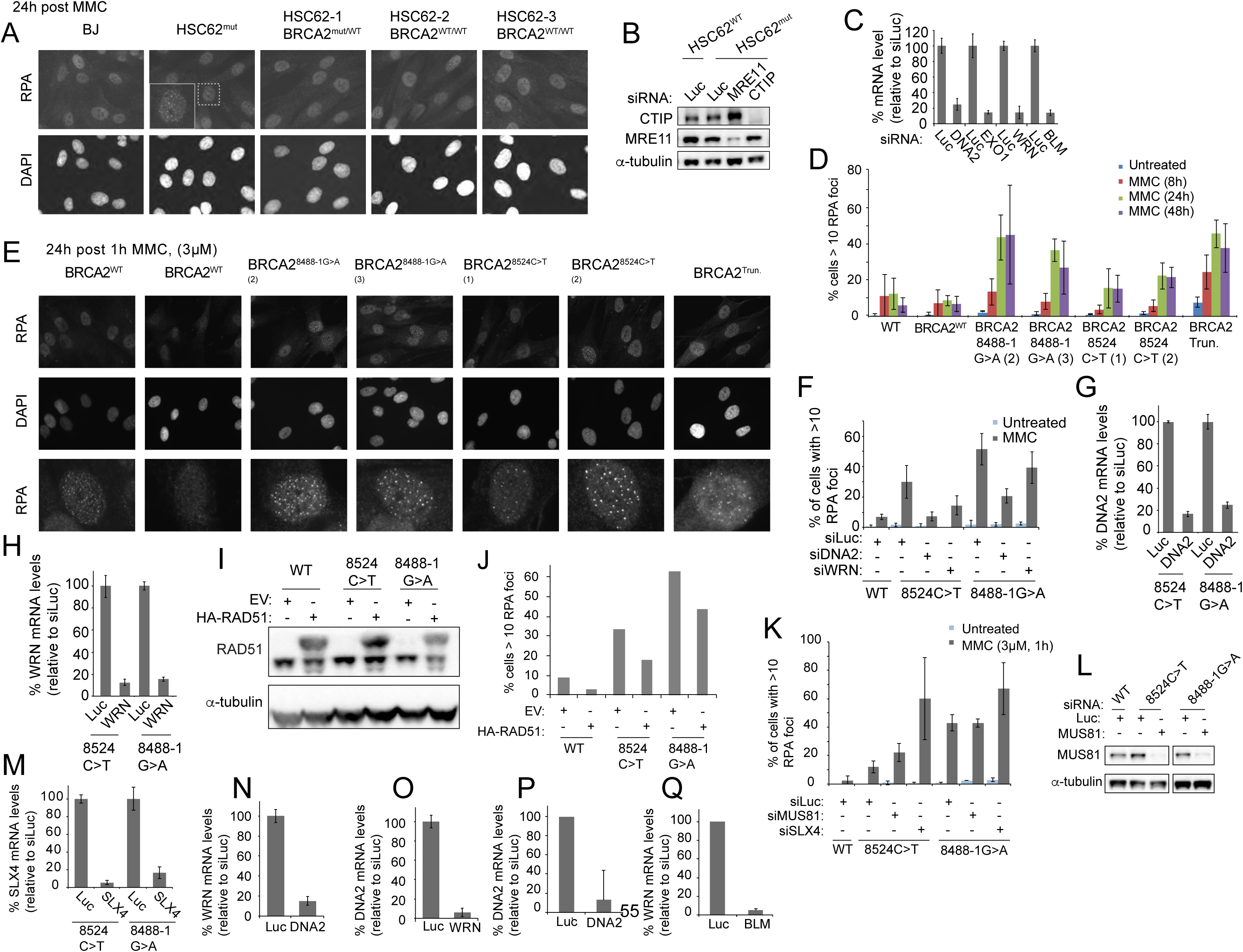
(A) Images of RPA foci in HSC62 uncorrected patient cell line (HSC62^mut^) compared to BJ WT fibroblast and CRISPR/Cas9 corrected wild type HSC62 (HSC62^mut/WT^ or HSC62^WT^) clones, 24h post 1h treatment with 3 μM MMC. Detected by immunofluorescence with anti-RPA32 antibody. Quantification in Figure 2G. (B)Immunoblot analysis of MRE11 and CTIP siRNA depletion for cells utilized in Figure 2H. (C) qRT-PCR of *DNA2, EXO1, WRN, and BLM* expression levels of cells in Figure 2H. Error bars are s.d. (D) Quantification of RPA foci in isogenic BJ fibroblasts clones 8h, 24h and 48h following 1h treatment with 3 μM MMC of BJ WT fibroblasts, BJ WT fibroblast (BRCA2^WT^), c.8488-1G>A BJ clones 2-3, c.8524C>T BJ clones 1-2, and a BJ *BRCA2* truncation mutant (BRCA2^Trun.^). Error bars indicate s.d. of three independent experiments (≥200 cells per experiment). (E) Representative images of RPA foci in isogenic BJ fibroblasts clones, 24h post 1h treatment with 3 uM MMC, detected by immunofluorescence with anti-RPA32 antibody. Third row images are individual cells enlarged to better demonstrate RPA foci. Quantified in Figure S3D. (F) Quantification of RPA foci in isogenic BJ fibroblasts clones 24h following 1h treatment with 3 μM MMC in cells depleted of DNA2 or WRN by siRNA. Error bars indicate s.d of two independent experiments. (G-H) qRT-PCR of *DNA2* and *WRN* expression levels of cells utilized in Figure 2I and Figure S3F. Error bars indicate s.d. (I) Immunoblot analysis of WT RAD51 overexpression in BRCA2^8524C>T^ *and* BRCA2^8488-1G>A^ BJ fibroblast cells used in Figure 2J-K and Figure S3J. (J) Quantification of RPA foci 24h following 1h treatment with 3 μM MMC in cells expressing WT RAD51 or EV control. Representative data of 3 independent experiments. (K) Quantification of RPA foci in isogenic BJ fibroblasts clones 24h following 1h treatment with 3 uM MMC in cells depleted of SLX4 or MUS81 by siRNA. Error bars indicate s.d of two independent experiments. (L) Immunoblot analysis of MUS81 depletion for cells utilized in Figure S3I. (M) qRT-PCR of *SLX4* expression levels of cells utilized in Figure S3I. Error bars indicate s.d. (N-O) qRT-PCR of *DNA2* and *WRN* expression levels of FA-A fibroblast cell lines utilized in Figure 3B. Error bars indicate s.d. (P-Q) qRT-PCR of *DNA2* and *BLM* expression levels of FA-A fibroblast cell lines utilized in Figure 3C. Error bars indicate s.d.

**Figure S4:**
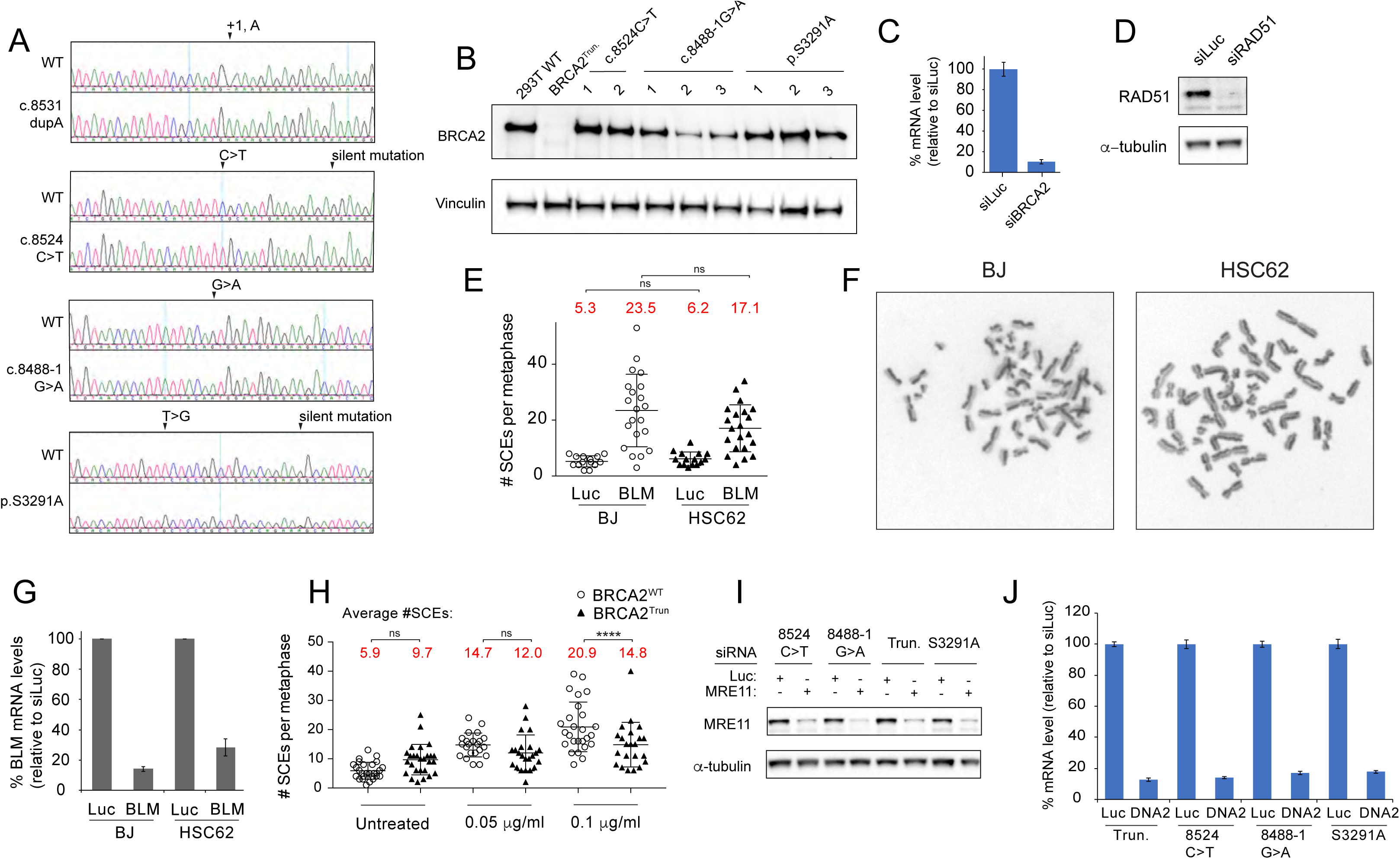
(A) Chromatograms of *BRCA2* CRISPR/Cas9 generated HEK293T clones aligned to WT. A frameshift Exon 20 BRCA2 mutant (BRCA2^Trun.^) was generated by homozygous single base pair insertion as a result of CRISPR/Cas9 targeting. *BRCA2* exon 20 variants, c.8524C>T and c.8488-1G>A, and Exon 27 p.S3291A (c.9871T>G) clones were generated by targeting the respective BRCA2 exon with CRISPR/Cas9 and a 100bp ssDNA template donor. Where applicable, silent mutations are indicated. Cell lines utilized in Figure 4A. (B) Immunoblot showing BRCA2 levels in WT HEK293T cells and *BRCA2* mutant HEK293T clones: c.8531dupA (*BRCA2^Trun^*), c.8524C>T (clones 1-2), c.8488-1G>A (clones 1-3), and p.S3291A (clones 1-3). Cells utilized in Figure 4A. (C) qRT-PCR of *BRCA2* expression levels of cells utilized in Figure 4A. Error bars are s.d. (D) Immunoblot of RAD51 knockdown for HEK293T cells used in Figure 4A. (E) SCE assay in BJ WT fibroblast and HSC62 fibroblast following depletion of BLM. (F) Representative images of SCEs in BJ WT fibroblast and HSC62 fibroblast metaphases. (G) qRT-PCR of *BLM* expression levels in cells described in E. Error bars indicate s.d. (H) Sister chromatid exchange (SCE) assay in BJ BRCA2^WT^ and BRCA2^Trun^ fibroblasts following treatment with MMC (0.05 μg/ml or 0.1 μg/ml). (I) Immunoblot analysis of MRE11 depletion for cells utilized in Figure 4D,F. (J) qRT-PCR of *DNA2* expression levels of cells utilized in Figure 4D,F. Error bars indicate s.d.

**Figure S5:**
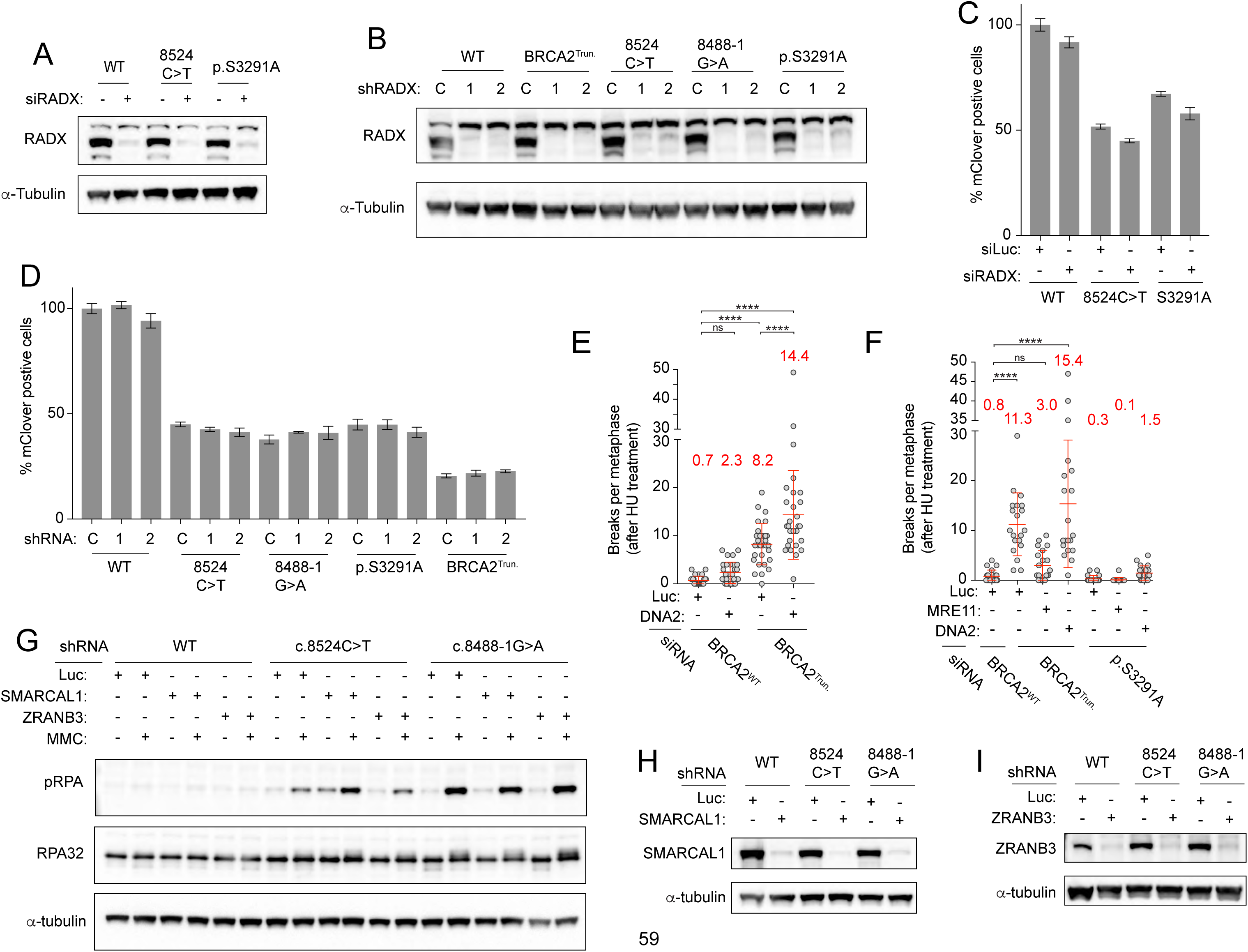
(A) Immunoblot analysis of RADX depletion by siRNA for cells utilized in Figure S5D. (B) Immunoblot analysis of RADX depletion by two shRNAs for cells utilized in Figure 4F. (C-D) Comparison of levels of mClover positive cells of HEK293T *BRCA2* mutants transfected with siRNAs or shRNAs targeting RADX or control (Luc or CONT). (E) Quantification of chromosome breaks in isogenic BJ BRCA2^WT^ and *BRCA2^Trun^*fibroblasts following 5h of 6 mM HU and released into colcemid. (F) Quantification of chromosome breaks in isogenic BJ *BRCA2^WT^* and *BRCA2^S3291A^* following 5h of 6 mM HU and released into colcemid. (G) Immunoblot analysis of RPA phosphorylation 24h post 1h treatment with 3 uM MMC. BRCA2^8524C>T^ *and* BRCA2^8488-1G>A^ BJ fibroblast cells were depleted of either SMARCAL1 or ZRANB3 by shRNA or transduced with shRNA control (Luc). (H-I) Immunoblot analysis of shRNA depletion of SMARCAL1 and ZRANB3 for cells utilized in Figure 5A-F and Figure S5E.

